# Genetic variations in G-Quadruplex forming sequences affect the transcription of human disease-related genes

**DOI:** 10.1101/2022.08.16.503999

**Authors:** Agustín Lorenzatti, Ernesto J. Piga, Mauro Gismondi, Andrés Binolfi, Ezequiel Margarit, Nora B. Calcaterra, Pablo Armas

## Abstract

Guanine-rich DNA strands can fold into non-canonical four-stranded secondary structures named G-quadruplexes (G4s). G4s folded in proximal promoter regions (PPR) are associated either with positive or negative transcriptional regulation. Given that single nucleotide variants (SNVs) affecting G4 folding (G4-Vars) may alter gene transcription, and that SNVs are associated with the human diseases’ onset, we undertook a comprehensive study of the G4-Vars genome-wide (G4-variome) to find disease-associated G4-Vars located into PPRs. We developed a bioinformatics strategy to find disease-related SNVs located into PPRs simultaneously overlapping with putative G4-forming sequences (PQSs). We studied five G4-Vars disturbing *in vitro* the folding and stability of the G4s located into PPRs, which had been formerly associated with sporadic Alzheimer’s disease (*GRIN2B*), a severe familiar coagulopathy (*F7*), atopic dermatitis (*CSF2*), myocardial infarction (*SIRT1*), and deafness (*LHFPL5*). Results obtained *in cellulo* for *GRIN2B* and *F7* suggest that the G4 disruption due to the identified G4-Vars affect the transcription and are responsible for the mentioned diseases. Collectively, data suggest that G4-Vars may account for the different susceptibilities to human genetic diseases’ onset, and could be novel targets for diagnosis and drug design in precision medicine.

## INTRODUCTION

Under certain conditions, specific DNA sequences do not adopt the canonical B-structure, but instead form non-B (non-canonical) DNA secondary structures such as cruciform, hairpin, i-motif, Z-DNA, and G-quadruplex (G4) (1), among others. G4s are dynamic structures formed by the self-folding of G-rich single-stranded DNA exposed during replication and transcription as a consequence of the negative supercoiling associated with these processes (2). Putative G4-forming sequences (PQSs) have been identified within the genomes of all kingdoms of life (3–5) and even in viral genomes (6), thus suggesting an important role of these structures throughout evolution.

The core structure of a G4 consists of a planar arrangement of four guanine residues bound by Hoogsteen base pairing conforming G-quartets or G-tetrads. The stacking of at least two G-tetrads establishes the G4, which is in turn stabilised by the coordination of monovalent cations, mainly by K^+^. The widely accepted consensus PQS is G_≥3_ N_1–7_ G_≥3_ N_1–7_ G_≥3_ N_1–7_ G_≥3_, wherein G-tracks are connected by loops of varying length and nucleotide (N) composition (7). However, non-canonical G4s can also form within sequences that have longer loops or less than three guanines per repeat (5, 8). The numbers of guanines per G-track and the length of the loops influence the stability of the G4 structure (9).

G4s were initially described in telomeric DNA sequences (10); but later, a significant number of studies have shown that G4s are important regulators of multiple cellular processes, such as replication, transcription, and genome maintenance (2, 11). Although the relevance of G4 structures in living cells was questioned in the past, accumulating experimental data now support the existence and importance of these structures in living cells and organisms (5, 12). The role of G4s in cancer is an area of intense study (13). G4s are found in telomeric DNA (14–16) and telomeric non-coding RNA (17), and play key roles in replication and genome instability (18, 19). Besides, most promoters of oncogenes harbour more G4 motifs than promoters of regulatory or tumour suppressor genes (20). Apart from the exhaustive studies that are being carried out to elucidate the role of G4 in the development of cancer, the exploration of the role of G4s in the context of other human diseases has grown-up remarkably in the last years (21).

Advances in the human genome sequencing during the last decades have shown sequence variations among individuals. Most of these variations consist of short indels and single-nucleotide variants (SNVs), which occur on average every 4–5 base pairs (22). Genome-wide association studies (GWAS) have identified more than 200.000 SNVs loci associated with several human diseases or traits. The majority of these SNVs are located in non-coding regions, including intergenic and intronic regions, and could affect gene expression by overlapping with transcription or translation regulatory elements (23). Therefore, one main challenge in human genetics is to understand the biological mechanisms by which SNVs influence phenotypes, including disease risk (24). The existence of 5 million gains/losses or structural conversions of G4s generated by the presence of SNVs has been recently reported. Of these, 3.4 million are within genes, preferentially enriched near the transcription start site (TSS) and overlapping with transcription factor binding sites and enhancers (25). Bearing in mind that G4s and the transcription factors associated with them cooperate to specify differential transcriptional programs (26), gains/losses or structural conversions of G4s generated by the presence of SNVs could not only alter a G4 structure but also influence gene expression among individuals (25, 27, 28).

A couple of works have reported a connection between disease risk and the gain/loss of G4s caused by SNVs. One of them reports that an SNV stabilising a G4 in the first intron of the calcium channel gamma subunit 8 gene (*CACNG8*) suppresses gene expression thus increasing the susceptibility to antisocial personality disorder (ASPD) (29). In the other, it was shown that the SNV responsible for the decrease in the transcription of the gene encoding placental anticoagulant protein annexin A5 (*ANXA5*) diminishes the potential for G4 formation *in vitro* and *in vivo*, suggesting an association between loss of the G4 and obstetric complications (30). Thus, we wondered whether gains/losses or structural conversions of G4s caused by disease-associated SNVs located in the proximal promoter regions (PPRs) of human genes influence their transcriptional expression. If so, not only the presence of a G4 but also its structural variants could impact on disease onset or susceptibility.

Here we report a comprehensive bioinformatics analysis performed by searching disease-related SNVs within PQSs located into the PPRs of human genes. We found novel PQSs that were characterised *in vitro* as being able to fold as G4s. In addition, we identified five SNVs associated with diseases that either promote or impair *in vitro* the formation and stability of G4s located into the PPRs of *GRIN2B, SIRT1, CSF2, F7*, and *LHFPL5* genes. Importantly, we gathered experimental data showing that SNVs present into the PPRs of *GRIN2B* and *F7*, previously reported to be responsible for altering the transcription of these genes and causing the related diseases (31, 32), affect the transcription of a reporter gene in cultured cells. Taken together, the data suggest that SNVs within G4s contribute to the onset or susceptibility of human pathologies.

## MATERIALS AND METHODS

### Obtaining human genomic variants and flanking sequences

Human Short Variants (HSV, including SNVs and indels excluding flagged variants) corresponding to HGMD-PUBLIC, ClinVar, and dbSNP databases, and Human Somatic Short Variants (HSSV, including SNVs and indels excluding flagged variants) corresponding to dbSNP, ClinVar, and COSMIC databases were downloaded from Ensembl Variation database (human reference genome GRCh38.p12; releases 92 and 93; https://www.ensembl.org/index.html) (33) using the Biomart tool. HSV and HSSV datasets were filtered to obtain disease-associated variants located into PPRs, defined here as the region spanning 1,000 bp upstream from the TSSs reported in Ensembl. The 50 bp up and downstream sequences flanking the variant positions were downloaded as multi-fasta files. In this work, and according to Ensembl, the “ancestral allele” (AA) was defined as the allele found in closely related species and is thought to reflect the allele present at the time of speciation. On the other hand, “variation allele” (VA) refers, in this work, to variation sequences derived from the ancestral ones. The nucleobases of both kinds of alleles (the AA and VA) were included as headers for every downloaded variant-flanking sequence. Separated multi-fasta sequence files were generated with a custom Perl script (Supplementary File S1), thus allowing the specific substitution of the variant position with the nucleobase corresponding to either the AA or VA, and the generation of two new multi-fasta files containing the substituted sequences; one of these new files represents AA-sequences and the other one the respective VA-sequences (Figure 1A). Since only the AA was reported for COSMIC and HGMD-PUBLIC variants, for the AA-sequences obtained from these databases, we generated the other three possible VA by replacing the AA-informed nucleobase with the other three possible nucleobases.

**Figure 1.**
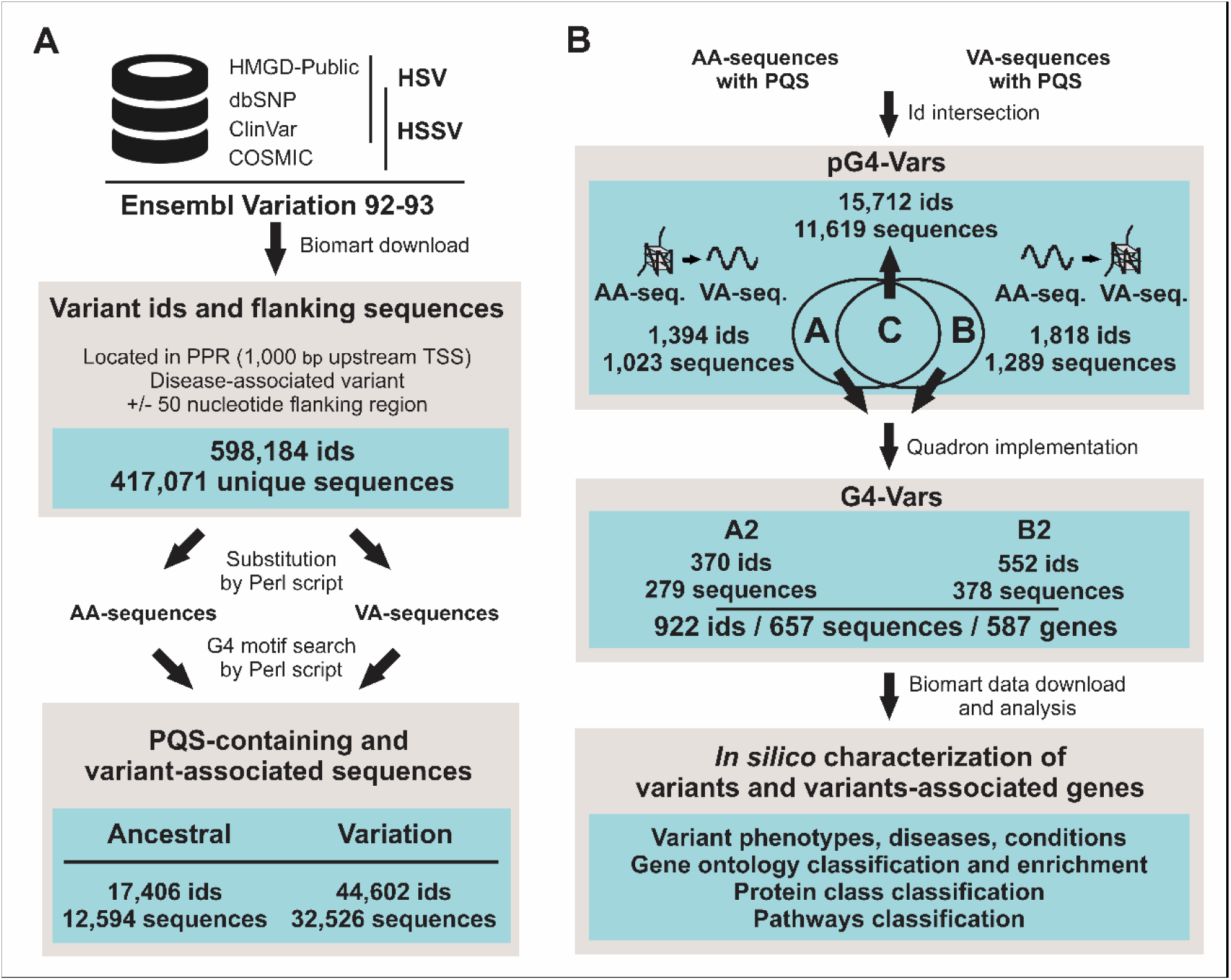
Bioinformatics pipeline towards the identification of disease-related G4-Vars. **(A)** Variant-associated sequences were downloaded from the Ensembl Variation database. A custom Perl script was designed to generate AA- and VA-sequences. Another custom Perl script was used to search for PQSs on both strands of AA- or VA-sequences. **(B)** Intersections between id lists of AA- and VA-sequences containing PQSs allowed identifying groups A and B of pG4-Vars. Quadron implementation yielded sub-groups A2 and B2 of G4-Vars (from groups A and B, respectively). Finally, variant phenotypes and variant-associated genes were *in silico* characterised according to Gene Ontology (GO) terms, Panther database annotations, and associated human diseases or conditions. Numbers of ids, sequences and genes obtained in each step of the pipeline (grey boxes) are shown (light blue boxes).

### PQSs finding, variant id intersections, and assessment of G4 formation propensity

A Perl script (Supplementary File S2) was generated to search for PQSs in both strands of the downloaded DNA sequences. We used the extended PQS definition consisting of four tracks of 3 to 7 guanines (G) (or cytosines (C) to consider the complementary strand) separated by three loops of 1 to 12 nucleotides (A, C, G, or T), according to Sahakyan et al. (34). The script reports the results in a tabular output file with the variant identification numbers (ids) contained in PQSs, the found PQSs, and the number of PQSs found per sequence. PQSs searches were performed on multi-fasta files generated for AA- and VA-sequences.

Variants occurring within PQSs (hereafter named as pG4-Vars) were analysed to identify those ones promoting or disrupting PQSs occurrence. Variant ids list intersections were performed by using Venny software (https://bioinfogp.cnb.csic.es/tools/venny/). Quadron package (http://quadron.atgcdynamics.org/) (34) was implemented on multi-fasta files containing the variant-flanking sequences (150 nucleotides both upstream and downstream, as required by the software) of the pG4-Vars downloaded from Ensembl. pG4-Vars within PQSs with Quadron Q parameter ≥ 19 were named here as G4-Vars.

### Oligonucleotides

The oligonucleotides representing the sequences studied in this work contained the complete PQSs flanked both at 5’ and 3’ ends by five additional nucleotides corresponding to the reference sequences informed in the Ensembl genome browser. For each case, mutations in PQSs were designed based on G-tracks disruption for impending G4 formation and were tested using the QGRS Mapper software (https://bioinformatics.ramapo.edu/QGRS/index.php) (35). Synthetic single-stranded oligodeoxyribonucleotides (Supplementary Table S1) were purchased from Macrogen Inc. with cartridge purification, dissolved in bi-distilled water and stored at −20°C until use. Concentrations were determined by spectrophotometry using extinction coefficients provided by the manufacturer.

### Circular dichroism (CD) spectroscopy

Intramolecular G4s were folded by dissolving 2 μM oligodeoxyribonucleotides in 10 mM Tris-HCl pH 7.5 and different KCl concentrations, as indicated in each figure, heating for 5 min at 95°C and slowly cooling to 20°C for at least 2 h. CD spectra were recorded at 20°C covering a wavelength range of 220–320 nm with a Jasco J-1500 spectropolarimeter (10 mm quartz cell, 1 nm band width, 1 nm data pitch, 100 nm/min scanning speed, 1 s response time). The recorded spectra represent a smoothed average of four scans, zero-corrected at 320 nm. The absorbance of the buffer was subtracted from the recorded spectra.

### 1D ^1^H Nuclear magnetic resonance (NMR)

Spectra were acquired at 20°C on a 700MHz Bruker Avance III spectrometer equipped with a triple resonance inverse NMR probe (5mm 1H/D-13C/15N TXI). Samples containing 50 µM oligodeoxyribonucleotides folded as G4 (as described for CD spectroscopy) in the presence of KCl, as indicated in each figure, were loaded into 5 mm Shigemi tubes. 1D ^1^H NMR spectra were registered using a pulse sequence with excitation sculpting (zgesgp) for water suppression (36). 8K points, 1K scans, a recycling delay of 1.4 sec and a sweep width of 18 ppm were used. Experimental time for each NMR spectrum was 30 min. Processing was done using an exponential window function multiplication with a line broadening of 10 Hz and baseline correction. Topspin 3.7 software (Bruker, Biospin) was used for acquisitions, processing, and analysis of the NMR spectra.

### qPCR Stop Assays (qPSA)

qPSA were performed as reported elsewhere (37), with some modifications. Briefly, each reaction tube (10 μL) consisted of 0.5 X SYBR Green, 500 nM of each PCR primer (Supplementary Table S1), 5 pM of templates oligodeoxyribonucleotides (AA, VA and M, Supplementary Table S1), 0.5 units of Platinum *Taq* DNA polymerase (Invitrogen), the PCR reaction buffer provided by the manufacturer, 2.5 mM of MgCl_2_ and 200 µM of each dNTP. Reactions were performed in a Realplex 4 Thermocycler (Eppendorf) using the following program: 95°C for 3 min (1 cycle) followed by a 2-step reaction of 95°C for 5 s and 60°C for 20 s (40 cycles), and finally a 20 min melting curve. Three technical replicates were performed for each experimental condition. Three independent qPSAs were done for each case. The validity of the qPCR data was assured by following the MIQE guidelines (38).To express the relative DNA amplification 2^-ΔCq^ values were determined, where ΔCq was defined as threshold cycle for either VA or M templates, minus threshold cycle of AA template. Data were statistically analysed by the T-Student test.

### Plasmids constructions

Duplex DNAs were generated by annealing oligodeoxiribonucleotides containing AA, VA or Mutated (M)-sequences with the corresponding complementary strands (Supplementary Table S1) and then cloned using SmaI restriction site upstream the basal SV40 promoter into pGL3-promoter vector plasmid (Promega), in the same strand (coding or template) as found in the genome, as previously described (39). Plasmids were sequenced and used for cell culture transfections as described below.

### Cell culture and transfection

HEK-293 cells were grown in Dulbecco’s Modified Eagle Medium (DMEM) and supplied with 10% foetal bovine serum (FBS), in a humidified cell incubator with an atmosphere of 5% CO_2_ at 37°C. Cells were seeded at a density of 3 × 10^6^ cells per 35 mm dish 24 h prior to transfection in DMEM containing 10% FBS. Cells were transfected using calcium phosphate method (40). Briefly, 250 ng of the pGL3-promoter vector (or the constructions derived from it) and 125 ng of the pCMV-β-gal plasmid (Promega) were used for transfection. The transfection medium was replaced by fresh medium 5 hours after transfection, and cells were grown as indicated above for 48 hours post-transfection (hpt).

### Luciferase reporter assays (LRAs)

LRAs were performed as previously described (39) with few modifications. At 48 hpt, cells were collected, lysated and firefly luciferase (FL) activities were measured using Luciferase Reporter Assay System (Promega, USA). FL activities were normalised to β-gal activities and expressed as the ratio of FL/βgal. Values determined for each construct were relativized to those for AA constructs for each PQS in study. The normalised ratio was obtained from three independent biological replicates for each experiment, and experiments were repeated three times. Data were statistically analysed by the T-Student test.

## RESULTS

### Obtaining a novel dataset of disease-related G4-Vars within PPRs

Short genetic variants may cause the gain, loss, and/or change in folding propensity of G4s that modulate transcriptional gene expression (25). Therefore, it is a fundamental issue to address the physiological function and pathological implications that such short genetic variations may represent. In view of this, we developed a bioinformatic pipeline (Figure 1) with the purpose of identifying disease-related genetic variants either promoting or disrupting the formation of G4s. We downloaded the disease-related HSV and HSSV contained into PPRs (defined as 1,000 bp upstream the TSS), along with their 50 bp-flanking sequences reported in the databases included in Ensembl Variation database (Figure 1A and Supplementary Table S2). Among the 598,184 downloaded sequences, 417,071 were unique sequences, as some of them were repeated due to being represented by different variant ids. From the 417,071 sequences, we generated two new sets of sequences, one corresponding to the ancestral allele (AA)-sequences and another to the variation allele (VA)-sequences. Both sets of sequences were used as inputs for a Perl script designed to search for an extended PQS definition consisting of four G-tracks of at least 3 Gs interspersed with extended loops of 1 to 12 nucleotides (G_≥3_ N_1–12_ G_≥3_ N_1–12_ G_≥3_ N_1–12_ G_≥3_), which accounts for 2/3 of the experimentally observed G4s (34) (Figure 1A and Supplementary Tables S3 and S4).

We identified 17,106 ids that resulted in AA-sequences containing PQSs corresponding to 12,594 unique sequences. On the other hand, we identified 44,602 ids that resulted in VA-sequences containing PQSs corresponding to 32,526 unique sequences (Figure 1A). For a better understanding, hereafter SNVs occurring within PQSs are named as pG4-Vars. The intersection between both id lists resulted in three groups. Group A (Supplementary Table S5 and Supplementary File S3) contains ids with PQSs only in the AA-sequences, group B (Supplementary Table S6 and Supplementary File S4) contains ids with PQSs only in the VA-sequences (Figure 1B), and group C, which contains the majority of the ids, includes ids presenting PQSs both in the AA- and VA-sequences. In the case of group C, the different alleles did not generate PQSs gains/losses. One possible explanation is that SNVs modify the sequence of the loops, the flanking regions or even the G-tracks, but do not affect the PQS consensus. Therefore, sequences belonging to group C were not further assessed in this study.

The 2,312 unique sequences contained in groups A and B (1,023 in group A + 1,289 in group B) were further characterised by the implementation of Quadron algorithm (34) in order to classify them according to the propensity of G4 formation. pG4-Vars within PQSs displaying Quadron scores higher than 19 (hereafter named as G4-Vars) were grouped in two new sub-groups, A2 and B2 (Figure 1B). Sub-group A2 comprises sequences containing PQSs displaying high propensity to fold as G4 only in the AA-sequences, while sub-group B2 consists of sequences containing PQSs with high propensity to fold as G4 only in the VA-sequences. Overall, 922 ids corresponding to 657 unique sequences were identified as disease-related G4-Vars located into PPRs (Figure 1B and Supplementary Table S7). Noteworthy, only four ids from the 370 found in sub-group A2 and only one from the 552 found in sub-group B2 are classified by Ensembl as variants located into PPRs (described as “upstream gene variant” by Ensembl) (Supplementary Figure S1). Instead, most variants are classified by Ensembl with calculated variant consequences overlapping with transcripts (5’ UTR, coding sequences, 3’ UTR, and introns). The low number of G4-Vars found located into PPRs may be explained at least by three possible scenarios. First, Ensembl reports all the possible transcripts for each gene, which are expressed in different tissues or developmental stages, but may not be the most abundant or representative transcripts. So, in our analyses, many G4-Vars located upstream of a TSS of a gene with several alternative TSSs may be classified by Ensembl as located within longer transcripts. Second, some G4-Vars may overlap with the transcription unit of another gene, being the variant consequence within the transcription unit probably prevalent over the upstream gene variant consequence. Third, an important number of G4-Vars are classified as overlapping with non-coding transcripts, thus they could be located within PPRs overlapping with non-coding transcripts, being the non-coding transcripts variant consequence prevalent over the upstream gene variant consequence. Therefore, it would be expected that more than the five G4-Vars described here as “upstream gene variant” are relevant for transcriptional control. It is worth mentioning that the SNVs located both into the PPR and simultaneously downstream the TSSs of an alternative transcript (within the transcriptional unit) may represent a complication in the analysis of consequences on gene expression due to putative effects of the G4-Vars on both transcriptional and post-transcriptional regulation of different transcripts for the same gene.

The variant ids from the sub-groups A2 and B2 (72 corresponded to the HSV and 850 to the HSSV datasets) were assessed for variant-associated phenotypes in the Ensembl Variation database. We found 833 ids (out of 922) related to a variety of pathologies, mainly different kinds of tumours (Supplementary Figure S2A). In addition, 922 variant ids were associated with 587 unique genes (according to the Ensembl Genes Database), which were characterised by considering gene ontology (GO) terms (Supplementary Figure S2B), Panther protein class (Supplementary Figure S2C), and pathway (Supplementary Figure S2D). Additionally, a GO enrichment study performed over the entire set of 587 genes showed that biological process terms related to development at different levels are prevalent (Supplementary Figure S2E and Supplementary Table S8).

### Selection of G4-Vars for experimental analyses

We focused the analyses on five G4-Vars described as “upstream gene variant” located into PPRs and upstream the TSSs of the most biologically relevant transcripts reported in Ensembl for the *GRIN2B, SIRT1, CSF2, F7*, and *LHFPL5* genes. *GRIN2B* encodes the glutamate ionotropic receptor NMDA type subunit 2B, a subunit of the N-methyl-D-aspartate (NMDA) receptor ion channels. Naturally occurring mutations within this gene are associated with neurodevelopmental disorders (41). The SNV causing the G4-Var identified here (Table 1 and Supplementary Figure S3A) increases *GRIN2B* transcriptional expression and was associated with reduced risk of sporadic Alzheimer’s disease (SAD) (31). *F7* encodes the Factor VII (FVII) protein, the serine-protease that triggers blood coagulation. A reduced plasma level of FVII results in bleeding diathesis of variable severity. The SNV causing the G4-Var identified here (Table 1 and Supplementary Figure S3B) causes a reduction in *F7* gene transcriptional expression and is responsible for a severe bleeding familiar disorder due to factor VII deficiency (32). *CSF2* encodes the cytokine colony stimulating factor 2, which has been related with pulmonary alveolar proteinosis and mucositis. The SNV causing the G4-Var identified here (Table 1 and Supplementary Figure S3C) is associated with reduced severity in atopic dermatitis (42). *SIRT1* encodes SIRTUIN 1, a member of a family of NAD-dependent class III deacetylases that modulate chromatin function and has been connected to many cellular processes such as cell cycle, response to DNA damage, metabolism, apoptosis, and autophagy. *SIRT1* is implicated in different human diseases and the SNV causing the G4-Var identified here (Table 1 and Supplementary Figure S3D) is associated with increased risk of myocardial infarction (43). *LHFPL5* encodes LHFPL tetraspan subfamily member 5, a member of the lipoma HMGIC fusion partner (LHFP) family, a subset of the superfamily of tetraspan transmembrane proteins. Mutations in this gene result in deafness, and it is proposed to function in the inner ear as a component of the hair cell’s mechanotransduction machinery (44). The SNV causing the G4-Var identified here (Table 1 and Supplementary Figure S3F) is associated with deafness (45).

**Table 1.**
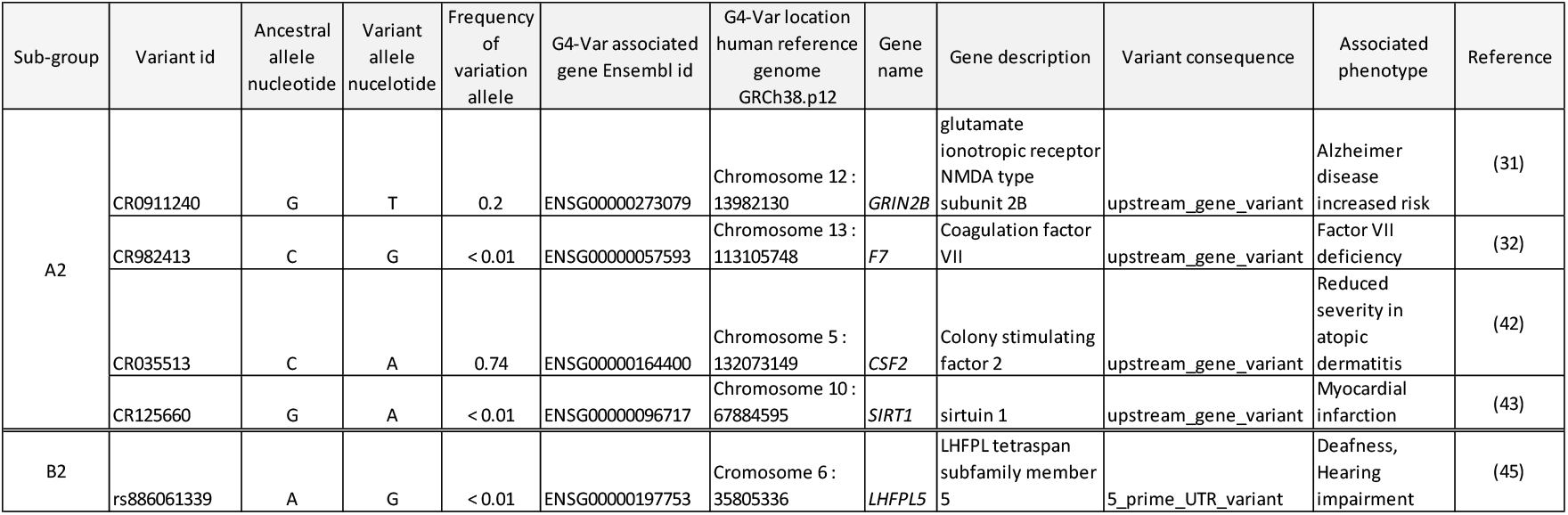
Summary of the data for the selected G4-Vars from sub-groups A2 and B2.

| Sub-group | Variant id | Ancestral allele nucleotide | Variant allele nucleotide | Frequency of variation allele | G4-Var associated gene Ensembl id | G4-Var location human reference genome GRCh38.p12 | Gene name | Gene description | Variant consequence | Associated phenotype | Reference |
| --- | --- | --- | --- | --- | --- | --- | --- | --- | --- | --- | --- |
| A2 | CR0911240 | G | T | 0.2 | ENSG00000273079 | Chromosome 12 : 13982130 | *GRIN2B* | glutamate ionotropic receptor NMDA type subunit 2B | upstream_gene_variant | Alzheimer disease increased risk | (31) |
|  | CR982413 | C | G | < 0.01 | ENSG00000057593 | Chromosome 13 : 113105748 | *F7* | Coagulation factor VII | upstream_gene_variant | Factor VII deficiency | (32) |
|  | CR035513 | C | A | 0.74 | ENSG00000164400 | Chromosome 5 : 132073149 | *CSF2* | Colony stimulating factor 2 | upstream_gene_variant | Reduced severity in atopic dermatitis | (42) |
|  | CR125660 | G | A | < 0.01 | ENSG00000096717 | Chromosome 10 : 67884595 | *SIRT1* | sirtuin 1 | upstream_gene_variant | Myocardial infarction | (43) |
| B2 | rs886061339 | A | G | < 0.01 | ENSG00000197753 | Chromosome 6 : 35805336 | *LHFPL5* | LHFPL tetraspan subfamily member 5 | 5_prime_UTR_variant | Deafness, Hearing impairment | (45) |

*GRIN2B, SIRT1, CSF2* and *F7*, G4-Vars were identified as potentially disrupting the formation of G4s (sub-group A2) and are described in Ensembl as “upstream gene variant”. In the case of *LHFPL5*, it was identified as potentially promoting the formation of G4s (sub-group B2) and, although the G4-Var is classified in Ensembl as “5 prime UTR variant”, is also located upstream of the TSS of the most biologically relevant transcript (main functional isoform, most conserved, highly expressed, and that has the longest coding sequence) (Figure 2A, Table 1 and Supplementary Figure S3F).

**Figure 2.**
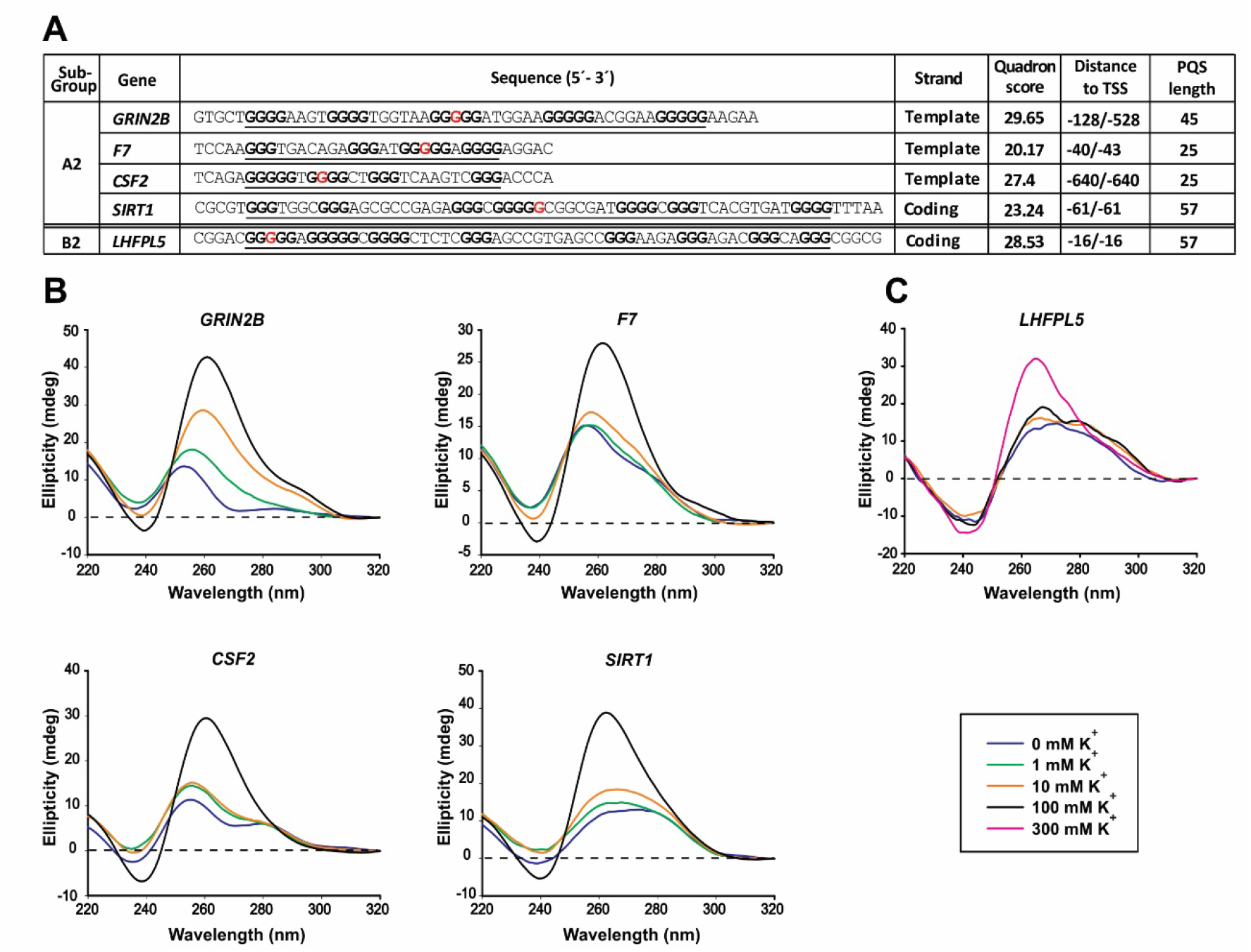
Evidence of *in vitro* G4 folding of PQSs containing the selected G4-Vars. **(A)** Main features of the sequences containing the PQSs including the five selected G4-Vars. Sequences from the sub-group A2 correspond to AA-sequences, while the sequence from the sub-group B2 correspond to VA-sequence. PQSs are underlined, G-tracks ≥ 3 are signalled in bold and the nucleobases involved in the G4-Vars are signalled in red. Distance to TSS informs both the distance to the most proximal downstream transcript and the distance to the most biologically relevant transcript according to Ensembl. **(B)** CD spectra obtained for the AA-sequences containing the PQSs of the four selected G4-Vars from the sub-group A2. **(C)** CD spectra obtained for the VA-sequence containing the PQS of the selected G4-Var from the sub-group B2. Oligonucleotides containing the PQSs were folded in the presence of increasing K^+^ concentrations, as indicated (right-bottom corner box).

G4-Vars are located within the most proximal 200 nucleotides upstream the TSS in the cases of *SIRT1, F7* and *LHFPL5*, or within the 500 nucleotides most distal of the PPRs upstream the TSS in the cases of *GRIN2B* and *CSF2*. Moreover, G4-Vars in *GRIN2B, SIRT1*, and *F7* are located on the template strand while G4-Vars in *CSF2* and *LHFPL5* on the coding strand in (Figure 2A and Supplementary Figure S3).

### *In vitro* analysis of G4-forming capability of PQSs containing G4-Vars

None of the PQSs containing the selected G4-Vars had been previously characterised as G4-forming sequences. Therefore, we performed dot-blots and CD spectra to assess whether G4 formation is feasible by synthetic single-stranded oligodeoxyribonucleotide sequences (Supplementary Table S1 and Figure 2A) containing each of the four PQSs from sub-group A2 or the PQS from sub-group B2. Dot-blots using BG4 antibody (46) show strong signals in the case of *GRIN2B* and *SIRT1*, mild signals in the case of *CSF2* and *F7* genes and no signal in the case of *LHFPL5* (Supplementary Figure S4). As expected, no signal was detected when assessing the corresponding mutated sequences (negative controls), in which G to A replacements disrupt G4 formation (Supplementary Table S1 and Supplementary Figure S4). In all cases, CD spectra show the typical pattern of peaks associated with parallel G4 structures, showing an increase of a positive peak around 264 nm and a negative peak around 240 nm in response to the presence of increasing K^+^ concentrations (Figure 2B-C). CD positive peaks reached the maximal intensities in the presence of 100 mM K^+^ (Figure 2B), except for the PQS of *LHFPL5*, which reached an evident CD spectrum of parallel G4 at 300 mM K^+^, probably indicating less G4 stability or formation propensity (Figure 2C). Noticeable, the five genes selected in this work were identified as containing observed G4 sequences (OQs) by a G4 high-throughput sequencing method to experimentally detect and map G4 structures in the human genome (G4-seq) (8). Also, the PPRs of these genes were identified as containing OQs (5). Moreover, a PQS was identified into the PPR of *SIRT1* in a genome-wide mapping of endogenous G4s by chromatin immunoprecipitation and high-throughput sequencing (G4 ChIP-seq) in the HaCaT cell line (47), indicating G4 formation within this PPR and probably an accessible open chromatin state linked to a high transcriptional activity of this gene. Taken together, data suggest that the PQSs containing the five selected G4-Vars are able to fold as G4 *in vitro*.

### *In vitro* analysis of the effect of G4-Vars on G4s formation

The effect of the G4-Vars on G4s formation was analysed by four different spectroscopic approaches (CD spectra, CD melting, TDS and 1D ^1^H NMR) and one biochemical method (qPSA). For the spectroscopic approaches, we used the synthetic single-stranded oligodeoxyribonucleotide representing the VA-sequences (for sub-group A2, *GRIN2B, SIRT1, CSF2* and *F7*) or the AA-sequence (for sub-group B2, *LHFPL5*). Additionally, sequences containing G to A site-specific mutations disrupting PQSs were used as negative controls (Supplementary Table S1 and Figures 3A, E, I and 4A, F). The VA-sequences from sub-group A2 display CD spectra with reduced intensities in the characteristic peaks of parallel G4s (mainly the positive peaks around 264 nm) when compared with the CD spectra for AA-sequences (Figures 3B, F and 4B, G), suggesting that the nucleotide changes reduce the G4s population likely by reducing their stability. A similar behaviour is observed for the AA-sequence from sub-group B2 (*LHFPL5*) when compared with the CD spectrum for the VA-sequence (Figures 3J). Regardless of the K^+^ concentration, mutated sequences show CD spectra with peaks not characteristic of G4 (Figures 3 and 4). G4 thermal stabilities calculated by CD melting show that melting temperatures (Tm) obtained for the VA-sequences from sub-group A2 are lower than those obtained for the AA-sequences, mainly for *CSF2* and *F7* (Supplementary Figure S5). On the contrary, the Tm obtained for the AA-sequence from sub-group B2 (*LHFPL5*) is lower than that obtained for the VA-sequence (Supplementary Figure S5). In agreement, TDS spectra for the AA-sequences from sub-group A2 and the VA-sequence from sub-group B2 show the typical G4 signature with two positive peaks around 243 and 273 nm and a negative peak at 295 nm (Supplementary Figure S6). Except for *SIRT1*, the VA-sequences from sub-group A2 and the AA-sequence from sub-group B2 show TDS spectral changes consistent with G4 structural changes. In all cases, the mutated sequences show TDS spectra not characteristic of G4. 1D ^1^H NMR showed a group of defined imino protons signals around 11–12 ppm for the five selected PQSs containing the AA-sequences in the case of PQSs from sub-group A2 or the VA-sequence in the case of the PQS from sub-group B2 (Figures 3C, G, K and 4C, H), confirming the presence of Hoogsteen bonds and G4 structures (48). All the VA-sequences from sub-group A2 and the AA-sequence from sub-group B2 display qualitative (Figures 3C, G, K and 4C, H) and quantitative (Supplementary Figure S7) differences in 1D ^1^H NMR spectra when compared with their corresponding AA- and VA-sequences. Indeed, although spectra show G4 signatures, the intensity of signals are lower, suggesting that the G4-Vars reduce the G4 population, likely by decreasing G4 stability. 1D ^1^H NMR spectra for the PQSs within the PPRs of *SIRT1, CSF2* and *LHFPL5* show signals around 13 ppm, indicating that these sequences may form structures with Watson-Crick base pairs at some extent (48) that may compete with G4s. As expected, the mutated sequences show spectra with no peaks indicating Hoogsteen bonds (i.e., no G4 structures).

**Figure 3.**
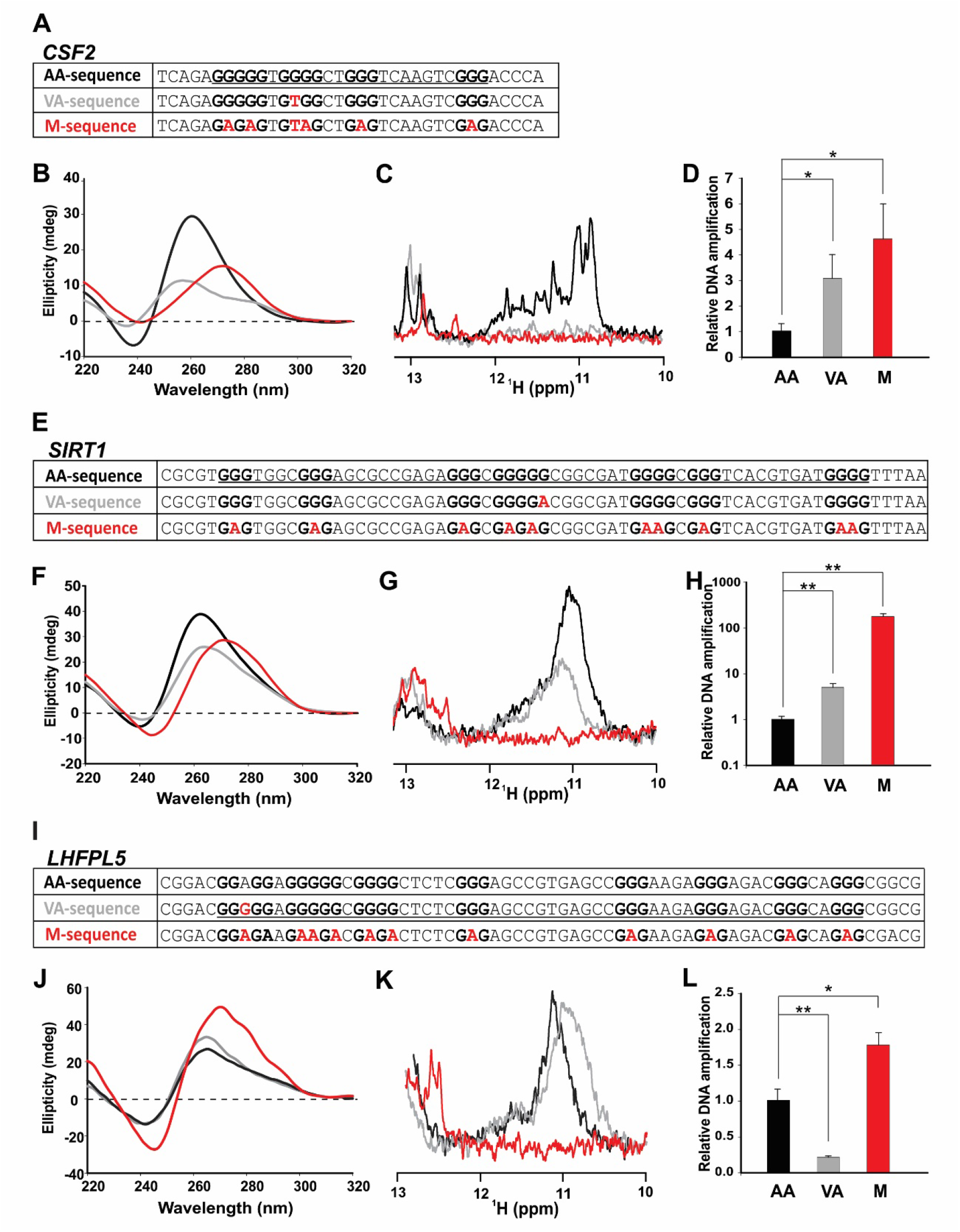
*In vitro* analysis of G4-Vars consequences in G4 formation by PQSs of *CSF2, SIRT1* (sub-group A2) and *LHFPL5* (sub-group B2). **(A), (E)** and **(I)** show tables with the AA-, VA-, and mutated (M)-sequences containing the PQSs of each selected G4-Var. PQSs are underlined, G-tracks ≥ 3 are signalled in bold and the nucleobases involved in the G4-Vars are signalled in red. **(B), (F)** and **(J)** show the CD spectra performed for oligonucleotides folded in the presence of 100 mM K^+^ (B and F) or 300 mM K^+^ (J). **(C), (G)** and **(K)** show the imino region of the 1D ^1^H NMR spectra obtained for each oligonucleotide sequence folded in the presence of the same K^+^ concentration used for CD. **(D), (H)** and **(L)** qPSAs results showing the relative DNA amplification. Results for AA-, VA-, and M-sequences are represented with black, grey, and red colours, respectively. *p < 0,05, **p< 0.01, T-Student test.

**Figure 4.**
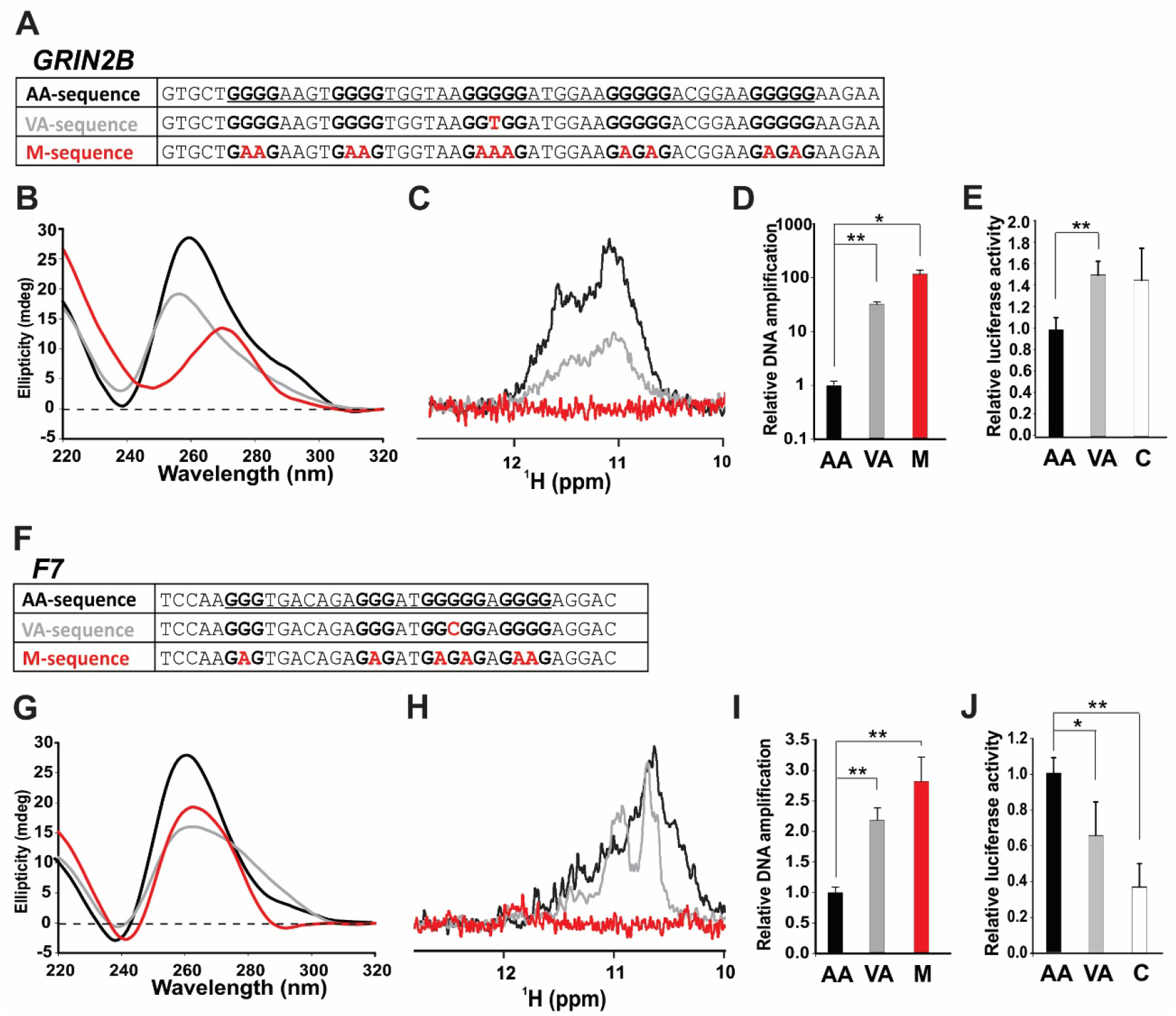
*In vitro* and *in cellulo* analysis of G4-Vars consequences in G4 formation by PQSs of *GRIN2B* and *F7* (sub-group A2). **(A)** and **(F)** show tables with the AA-, VA-, and M-sequences containing the PQSs of each selected G4-Var. PQSs are underlined, G-tracks ≥ 3 are signalled in bold and the nucleobases involved in the G4-Vars are signalled in red. **(B)** and **(G)** show the CD spectra performed for oligonucleotides folded in presence of 10 mM K^+^ (B) or 100 mM K^+^ (G). **(C)** and **(H)** show the imino region of the 1D ^1^H NMR spectra obtained for each oligonucleotide sequence folded in the presence of the same K^+^ concentration used for CD. **(D)** and **(I)** qPSAs results showing the relative DNA amplification. **(E)** and **(J)** LRAs were performed in HEK-293 cells transfected with pGL3-promoter vector containing AA- or VA-sequences for each PQS inserted upstream the basal promoter SV40, as well as with a control (C) with no insert. Bars represent luciferase activity (mean of three independent experiments) normalised to β-galactosidase activity and relativized to values obtained for the corresponding AA-sequence. Error bars correspond to standard deviation (SD). Results for control (C), AA-, VA-, and M-sequences are represented with white, black, grey and red colours, respectively. *p < 0.05, **p < 0.01, T-Student test.

The qPSA relies on the amplification of ssDNA templates consisting of a central PQS and flanking sequences for the annealing of primers. G4s formed in the ssDNA templates act as roadblocks to *Taq* polymerase, which stalls replication and leads to a decreased amplification efficiency by qPCR. Therefore, formation of G4s with higher stabilities will boost this effect and reduce the amplification efficiencies, while G4 destabilisation will increase the amplification efficiency (37). Figures 3D, H, L and 4D, I show that qPSA amplification efficiencies for the VA-sequences from sub-group A2 and the AA-sequence from sub-group B2 are higher than the qPSA amplification efficiencies for their AA-sequences and VA-sequence, respectively, confirming that G4-Vars disrupting PQSs reduce the G4s population likely by reducing their stabilities. In all cases, the mutated sequences show higher qPSA amplification efficiencies indicating no G4 formation and maximal amplification.

Pyridostatin (PDS) is a widely used G4 stabiliser molecule (5, 8, 49). CD spectra performed on sequences pre-incubated with PDS show that the sequences with disrupted PQSs are more responsive to increasing PDS concentrations than their counterparts containing PQSs (Supplementary Figure S8). qPSAs performed in the presence of PDS (Supplementary Figure S9) show that the VA-sequences for *GRIN2B, CSF2*, and *SIRT1* are more susceptible to be stabilised by PDS than the AA sequences. These results suggest that G4-stabilising ligands revert the effect of the G4-Vars on G4s stabilities.

Collectively, *in vitro* results suggest that the selected G4-Vars affect G4s formation and stabilities, which can be reverted by the presence of G4-specific ligands. These facts may account for functional effects in transcriptional gene expression control.

### *In cellulo* analysis of G4-Vars consequences on transcriptional regulation

The next key question was whether the G4-Vars have consequences on transcriptional activity. To address this, we focused on the two G4-Vars for which there exists bibliographic information about an effect on transcriptional expression. The SNV causing the G4-Var identified within the PPR of *GRIN2B* gene (CR0911240 retrieved from HGMD-PUBLIC) had been previously found in a systematic screening of the PPR of *GRIN2B* in the North Chinese population for detecting possible genetic variants genetically and functionally associated with sporadic Alzheimer’s disease (SAD). Genetic analysis further confirmed that homozygosis in AA (defined as mutant-type haplotype) increase the risk for SAD, even within subjects without APOE ε4 allele (considered a risk-factor for AD and for early age of disease onset). Importantly, this SNV leads to lower *GRIN2B* gene transcriptional expression (31). On the other hand, the SNV causing the G4-Var identified within the PPR of *F7* gene (CR982413 retrieved from HGMD-PUBLIC) had been previously identified as a mutation responsible for a severe bleeding familiar disorder (factor VII deficiency, hemarthrosis and chronic arthropathy) in a patient with homozygous VA (defined as mutant-type haplotype), and causing a reduction in *F7* gene transcriptional expression (32). Therefore, for both cases, we constructed luciferase reporter plasmids by cloning either the AA- or the VA-sequences upstream of the SV40 basal promoter into the pGL3 promoter vector. Each PQS was cloned in the same DNA-strand (template in both cases) in which they are found in the human genome. Plasmids were used for transient transfections in HEK-293 cells and the luciferase activity was measured as a transcriptional reporter (Figure 4E, J).

Luciferase activity obtained using the plasmid containing the *GRIN2B*-AA-sequence is significantly lower than that measured using the plasmid containing the *GRIN2B*-VA-sequence and for the empty pGL3-promoter vector, while the activity detected for the plasmid containing the *GRIN2B*-VA-sequence is similar to that observed for the empty pGL3-promoter vector. Results suggest that the *GRIN2B*-AA-sequence capable of folding as G4 has a repressive effect on transcription. The G4-Var may release the repression, probably by destabilising or loosening the G4 structure, thus reaching luciferase activities similar to those for the empty pGL3-promoter vector. Importantly, these results are in agreement with data reported by Jianga and Jia (31) who, using a fragment of the PPR of *GRIN2B* containing the AA-sequence, observed lower luciferase activity than that for the VA-sequence when transfected in human neuroblastoma (SH-SY5Y) and human epithelial carcinoma (HeLa) cell lines.

Luciferase activity obtained using the plasmid containing the *F7*-AA-sequence is significantly higher than that obtained using the plasmid containing the *F7*-VA-sequence, while the activity detected for the empty pGL3-promoter vector is even lower than that one observed for the plasmid containing the *F7*-VA-sequence. Results suggest that the *F7*-AA-sequence capable of folding as G4 may enhance transcription, since luciferase activities were significantly higher than those measured for the empty pGL3-promoter vector. Moreover, the nucleotide change found on the *F7*-VA-sequence reduces this effect, probably by destabilising or loosening the G4 structure. These results are in agreement with reporter gene assays performed using a region of the PPR of *F7* showing that the *F7*-VA-sequence displayed a 20-fold reduction in reporter gene expression compared with that for the *F7*-AA-sequence when transfected in human hepatoma (HepG2) cell line (32). In agreement, LRA experiments performed using a different DNA fragment within the PPR of *F7* showed that the *F7*-VA-sequence displayed lower luciferase activity than that for the *F7*-AA-sequence when transfected in HepG2 cell line (50).

Bibliographic data along with *in cellulo* results presented here suggest that the selected G4-Vars in the context of the PPRs cause changes in the G4s folding and stability and may account for changes in the expression of genes that could be relevant for the onset or susceptibility of human genetic diseases.

## DISCUSSION

Genetic variations throughout the genome found within PQSs and affecting G4s formation (G4-Vars) can be envisaged as the “G4-variome”. The contribution of the G4-variome to the functional diversity of the human genome has been brought to light during the last decade. Indeed, several studies focused on RNA G4s have shown that G4-Vars may perturb the translation, stability, and localization of mRNAs (51–54), and affect the microRNA biogenesis (55). The potential impact of the G4-Vars on DNA biology began to emerge by the identification G4-Vars in genome-wide studies (25, 27, 28, 56–58) and by the characterisation of the role of specific G4-Vars in the transcriptional regulation of particular genes (29, 30). A pioneer genome-wide analysis of SNVs in human PQSs was performed emphasising on G4-Vars occurrences both in genes and their regulatory sequences. Data suggested that disruptive variations in G-tracks of PQSs are less frequent than neutral variations in loops (56). Then, a merged analysis of genotype information, SNV data, and gene expression profiles assessed whether the difference in the expression of particular genes is associated with the G4-Vars. Bioinformatics and experimental results suggested both a relative selection bias against alteration of PQSs and a significant role of G4-Vars in gene expression variations among individuals (27). Based on these findings, G4-Vars were suggested as *cis* quantitative trait loci associated with expression (*cis*-eQTLs, or loci responsible for quantitative alteration in gene expression) capable of causing substantial alteration of promoters activities (28). A more recent work has reported the existence of more than 5 million pG4-Vars causing gains/losses or structural conversions of PQSs within the human genome, most of them enriched near the TSSs and mainly overlapping with transcription factor–binding sites (TFBSs) and enhancers. This finding envisaged important putative implications of the G4-variome on gene activity and positioned the G4-Vars as a novel category of targets for personalised health risk assessment and drug development (25). Moreover, it has been proposed that G4-Vars are also relevant for gene expression variations in organisms of commercial interest, such as barley (57) and cattle (58), thus potentially linking G4-Vars to traits of agricultural and livestock importance.

Although several works have pointed out a link between the G4-variome and the appearance of diseases, a comprehensive search intended for identifying pathogenic G4-Vars with possible consequences on gene transcription has not yet been carried out. In this work, we developed a novel strategy aimed at identifying and testing disease-associated G4-Vars located into PPRs likely affecting transcriptional gene activity. It should be noted that our original *in silico* pipeline is not restricted solely to the goal of this work, and can also be adapted to identify G4-Vars affecting other steps of gene expression or even other pathways in which the G4s are involved.

We have set high stringent criteria for the selection of the G4-Vars to be assessed, being aware that this stringency may have circumvented some biologically relevant G4-Vars. We restricted the analysis to those G4-Vars with high chances of impairing or favouring G4s formation due to disruption or promotion of a PQS (groups A and B); i. e., G4-Vars that mainly affect G-tracks. Moreover, we selected by Quadron algorithm (34) the PQSs with the highest propensity to form G4 (sub-groups A2 and B2) with the extended canonical consensus definition. This consensus includes 65% of the observed G4s by G4-Seq (34), but do not consider the non-canonical sequences (that contain bulges in the G-tracks or loops longer than 12 nucleotides), which would provide additional complexity to the G4-variome scenario. In addition, we left behind G4-Vars probably containing SNVs within the loops, the flanking regions, or even the G-tracks as long as they keep the presence of the PQSs. These G4-Vars represent the majority of the identified pG4-Vars (83 %, or 15,712 ids out of a total of 18,924) and, even conserving a canonical PQS, could alter the G4s stability and/or topology, thus accounting for changes in transcriptional levels of genes regulated by G4s. Therefore, we consider this work as a start point for discovering the implications of the G4-variome in the onset of human genetic diseases or even in the possible treatment, diagnosis, and/or prevention of these diseases.

The five assessed G4-Vars comprise a variety of cases including G4 disruption (four G4-Vars from sub-group A2) and promotion (one G4-Var from sub-group B2), with PQSs located on the template (three G4-Vars) and coding (two G4-Vars) strands in respect to transcription, and at different distances from TSSs within the PPRs. The five G4-Vars had been formerly associated with pathological phenotypes of diverse nature, including neurological and cardiac diseases, atopic dermatitis, coagulopathy and deafness, but had not been related with G4 structures. By means of a battery of biophysical and biochemical methods, we confirmed that these G4-Vars are located into PQSs able to fold as G4s and verified the expected effect (i.e., G4 disruption or promotion), thus pointing out the success and robustness of the *in silico* strategy. In the cases of G4-Vars found in *GRIN2B* and *F7* promoters, results gathered in cultured cells recapitulated reports by other groups describing the effect of these SNVs on transcription (31, 32, 50). Results suggest that the G4-Vars identified here change the G4 folding capability and account for changes in the expression of genes relevant for the establishment of a severe coagulopathy (in the case of *F7*) or for the predisposition for Alzheimer’s disease (in the case of *GRIN2B*).

Although the influence of G4 structures *per se* on transcription activity has been widely described (2, 59), the combined effects of DNA-binding proteins acting as transcription factors (TFs) together with G4 folding should also be considered, as G4s may act as binding hubs for many different TFs influencing transcription (26, 60). Indeed, the G4-Var in the *GRIN2B* PPR overlaps with a putative binding site for the zinc finger ras-responsive element binding protein (RREB), reported as a transcriptional repressor. Therefore, it was suggested that the AA-sequence allows the binding of RREB, thus repressing transcription, while the VA-sequence impairs RREB binding, thus promoting transcriptional activity (31). Regarding the G4-Var found in the *F7* PPR, the VA-sequence prevents the binding of SP1 and other nuclear proteins leading to transcriptional repression (32, 50). The link between PQSs and the occurrence of the SP1-binding sequences has been extensively characterised (61–63), and SP1 has been described as a G4 binding protein (26, 64–66). Alterations in G4 folding may adversely impact on SP1 binding affinity and, consequently, on its functions as a TF (66, 67). In line with this, a predictive analysis of TFBSs performed on AA- and VA-sequences for the five G4-Vars assessed here shows that nucleotide variations modify the predicted TFBSs for several TFs in both the AA- and the VA-sequences (Supplementary Table S9 and Supplementary File S5). Among the TFBS differentially predicted between AA- and VA-sequences, several are G4-binding proteins or TFs formerly described as able to bind to sequences overlapping or close to PQSs; e.g., MAZ, E2F1, p53 and WT1. Therefore, it is tempting to speculate that the binding of these TFs is modified by changes in G4s folding or stabilities caused by G4-Vars, thus impacting on transcriptional activity.

This work is a pioneer exploration of the G4-variome carried out focusing on the transcription of genes related to human diseases. The identification of G4-Vars related to human pathologies is not only useful for disease diagnosis, but may also indicate the relevance of a particular G4 structure and thus serve as a target for drug development and design of disease-specific treatments. The effect of G4-Vars promoting or disrupting G4 formation could be reverted by treatments with specific ligand molecules destabilising or stabilising an affected G4 structure, as suggested by our results with PDS, thus restoring the gene expression levels. Therefore, scientific efforts should be focused on deepening in the discovery and design of sequence-specific G4 ligands, thus contributing to the design of personalised therapeutic approaches.

## Supporting information

Graphical Abstract

## ABBREVIATIONS

G4,: G-quadruplex;
PQS,: putative G4-forming sequence;
SNV,: single-nucleotide variant;
TSS,: transcription start site;
PPR,: proximal promoter region;
HSV,: Human Short Variants;
HGMD-PUBLIC,: Public Human Gene Mutation Database;
dbSNP,: Single Nucleotide Polymorphism database;
HSSV,: Human Somatic Short Variants;
COSMIC,: Catalogue Of Somatic Mutations In Cancer;
AA,: ancestral allele;
VA,: variation allele;
pG4-Var,: genetic variant occurring within a PQS;
G4-Var,: genetic variant occurring within a sequence that forms a G4;
QGRS,: Quadruplex forming G-Rich Sequence;
CD,: circular dichroism;
NMR,: nuclear magnetic resonance;
qPSA,: qPCR Stop Assays;
qPCR,: real-time quantitative PCR;
HEK-293,: human embryonic kidney 293 cell line;
DMEM,: Dulbecco’s Modified Eagle Medium;
FBS,: foetal bovine serum;
hpt,: hours post-transfection;
LRA,: luciferase reporter assay;
FL,: Firefly luciferase;
β-gal,: β-galactosidase;
GO,: gene ontology;
SAD,: sporadic Alzheimer’s disease;
OQ,: observed G4 sequence;
Tm,: melting temperature;
PDS,: pyridostatin;
(TFBS),: transcription factor–binding site;
TF,: transcription factors.

## DATA AVAILABILITY

All data obtained and presented in this work are available in the manuscript and as Supplementary Tables and Files. Quantitative PCR comply with the MIQE Guidelines as detailed in Materials and Methods section. Synthetic oligonucleotides sequences and source are detailed in Materials and Methods section and in Supplementary Table 1. Custom scripts are available as Supplementary Files 1 and 2. Complete data obtained from bioinformatics approaches are available as Supplementary Tables 2 to 9 and Supplementary Files 3 to 5.

## FUNDING

This work was supported by Agencia Nacional de Promoción Científica y Tecnológica [grant numbers PICT 2016-0671 to N.B.C., PICT 2017-0976 and PICT 2019-1662 to P.A.], Consejo Nacional de Investigaciones Científicas y Técnicas [grant number 2015-0170 to N.B.C.] and Universidad Nacional de Rosario [grant number BIO573 to P.A.].

## ACKNOWLEDGEMENTS

We are thankful to Dolores Campos for excellent cell culture assistance.

## AUTHOR CONTRIBUTIONS

P.A. and N.B.C. performed the conceptualization and design of the work. M.G. and E.M. developed the bioinformatic pipeline to identify disease-related G4-Vars, and *in silico* characterised their related genes. E.J.P. obtained the variation databases and sequences data and A.L. performed the analysis of transcription factors binding sites. A.L. performed most experiments with the assistance of E.J.P. A.B. performed and analysed NMR spectroscopies. P.A. and N.B.C. were responsible for supervision, funding acquisition, project administration, and obtaining of resources. P.A. conducted the original draft writing and visualization, while P.A., A.L., E.J.P. and N.B.C. performed the major writing review and editing, assisted by M.G., E.M. and A.B. All authors have read and agreed to the published version of the manuscript.

## Notes

### Competing Interest Statement

The authors have declared no competing interest.

