## Supplementary Material for "Genetic variations in G-Quadruplex forming sequences affect the transcription of human disease-related genes"

### Content:


| | Page |
|---|---|
| Supplementary Tables Legends | 1 |
| Supplementary Files Description | 3 |
| Supplementary Figures | 4 |
| Supplementary Figure 1 | 4 |
| Supplementary Figure 2 | 6 |
| Supplementary Figure 3 | 8 |
| Supplementary Figure 4 | 12 |
| Supplementary Figure 5 | 13 |
| Supplementary Figure 6 | 14 |
| Supplementary Figure 7 | 15 |
| Supplementary Figure 8 | 16 |
| Supplementary Figure 9 | 17 |
| References for Supplementary Material | 18 |

### Supplementary Tables Legends

**Supplementary Table S1. Oligonucleotides used in this work.** Names and sequences of the synthetic single-stranded oligodeoxyribonucleotides used for different purposes are presented.

**Supplementary Table S2. Downloaded human variants.** Variants ids downloaded from Ensembl Variation databases releases 92 and 93 with their source datasets (HSV -Human Short Variants- or HSSV -Human Short Somatic Variants-) and original source databases (dbSNP, ClinVar, COSMIC, HGMD-PUBLIC). Variants were filtered considering distance to transcription start site (TSS) ≤ 1,000 bp and reported association with human diseases or conditions.

**Supplementary Table S3**. **PQS-containing AA-sequences.** Output results obtained after implementing the developed Perl script (Supplementary File S2) on AA-sequences (previously substituted with the AA nucleotide version) are shown. Different sets of sequences were analysed based on the source dataset and database. The number of PQS-containing variants of the set, variant ids, AA nucleotide, number of PQSs found for each variant-associated sequence, and the PQS motif sequence(s) are included.

**Supplementary Table S4**. **PQS-containing VA-sequences.** Output results obtained after implementing the developed Perl script (Supplementary File S2) on VA-sequences (previously substituted with the VA nucleotide version) are shown. Different sets of sequences were analysed based on the source dataset and database. The number of PQS-containing variants of the set, variant ids, VA nucleotide version, number of PQSs found for each variant-associated sequence, and the PQSs are included.

**Supplementary Table S5. pG4-Vars from group A: variant-associated sequences in which PQSs were only found in the AA-sequences.** Variant id intersection of PQS-containing AA-sequences with PQS-containing VA-sequences revealed the group A that includes those variant ids that are only present within the AA set. Table shows: variant id, AA nucleotide, PQSs number and sequence.

**Supplementary Table S6. pG4-Vars from group B: variant-associated sequences in which PQSs were only found in the VA-sequences.** Variant id intersection of PQS-containing AA-sequences with PQS-containing VA-sequences revealed the group B that includes those variant ids that are only present within the VA set. Table shows: variant id, VA nucleotide, PQSs number and sequence.

**Supplementary Table S7. G4-Vars: variant-associated sequences with a differential propensity to fold as G4 according to the presented variant allele.** Table shows: Variant id, AA or VA nucleotide(s), original Ensembl Variation dataset and database source, PQS start position, strand and length, Q value (Quadron prediction parameter), found PQS(s) by Quadron, G4-Var-associated gene id, name, description, and variant consequence. Data is structured in two datasets named A2 and B2, being A2 a sub-group of the previously mentioned group A, and B2 a sub-group of group B.

**Supplementary Table S8.** ***In silico*** **characterization of G4-Vars and G4-Vars-associated genes.** Table shows G4-Vars ids, G4-Vars-associated phenotypes (HSV- and HSSV-G4-Vars associated phenotypes, along with a count of them), G4-Vars-associated genes (including gene stable ID, gene description, chromosome, gene start and end, strand, gene name), Panther analysis (including gene ontology (A, B, C), protein class (D), and pathways (E)), and gene ontology term enrichment analysis.

**Supplementary Table S9. Analysis of predicted TFBSs in AA- and VA-sequences for the five selected G4-Vars.** Table shows TFBSs predicted only in AA-sequences or with better match (lower dissimilarity) in AA-sequences; TFBSs predicted only in VA-sequences or with better match (lower dissimilarity) with VA-sequences; and references for those TFs previously associated with G4s. Predictions were performed using the PROMO software (1, 2), see Supplementary File 5.

### Supplementary Files Description

**Supplementary File S1. Perl script for generating AA- and VA-sequences.** Perl script designed and implemented to use sequences downloaded from the databases and all their alleles as inputs to generate two types of multi-fasta files: one containing the AA sequences and the other containing the VA sequences. The sequences in these new files retain sequence identifiers from the different variation databases and contain the specific allele (AA or VA) that was used for the substitution.

**Supplementary File S2. Perl script for PQS-searching tool.** Perl script designed and implemented to identify and report PQS sequences as cis-elements within input variant-associated sequences in a multi-fasta file. The script searches PQSs of four tracks of 3 to 7 guanines (G) or cytosines (C) separated by three linkers (spacers) of 1 to 12 N nucleotides (N = A, C, G, or T). This algorithm allows the user to get a tabular output file with sequence identifiers, the number of PQS-containing sequences, the PQS found sequences and their number per sequence. An additional output file of the same program is a multi-fasta file with the group of PQS-containing sequences from the input file.

**Supplementary File S3**. **Group A sequences.** Multi-fasta file with group A AA-sequences representing those cases in which only the AA-sequences allow PQS finding.

**Supplementary File S4. Group B sequences.** Multi-fasta file with group B VA sequences representing those cases in which only the VA-sequences allow PQS finding.

**Supplementary File S5. Raw data of the prediction of TFBSs in AA- and VA-sequences for the five selected G4-Vars.** PROMO software (1, 2) was used in order to predict those TFs that could bind differentially to both AA- and VA-sequences for the five cases in study. A summary of this analysis is shown in Supplementary Table S9.

### Supplementary Figures

**A**

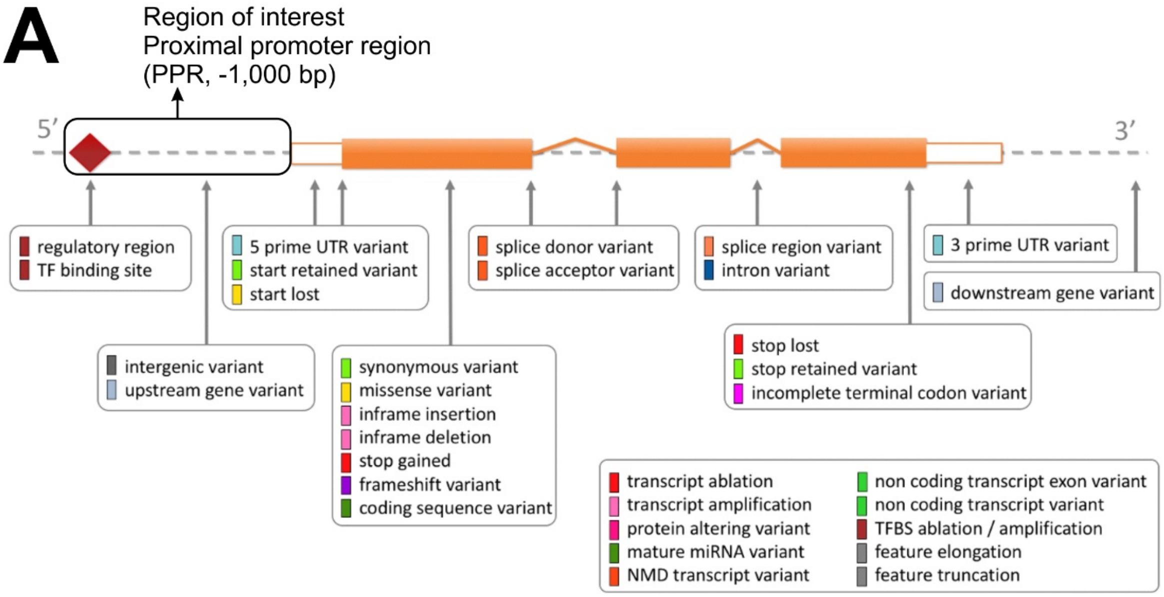


**B**

| Variant consequence | A2_G4-Vars | B2_G4-Vars | Total (A2+B2) |
|---|---|---|---|
| **upstream_gene_variant** | 4 | 1 | 5 |
| | | | 0 |
| **5_prime_UTR_variant** | 42 | 107 | 149 |
| start_retained_variant | 0 | 1 | 1 |
| start_lost | 0 | 0 | 0 |
| | | | 0 |
| **coding_sequence_variant** | 289 | 496 | 785 |
| synonymous_variant | 6 | 0 | 6 |
| missense_variant | 12 | 5 | 17 |
| inframe_insertion | 0 | 0 | 0 |
| inframe_deletion | 1 | 0 | 1 |
| stop_gained | 1 | 0 | 1 |
| frameshift_variant | 2 | 2 | 4 |
| stop_lost | 0 | 0 | 0 |
| stop_retained_variant | 0 | 0 | 0 |
| incomplete_terminal_codon_variant | 0 | 1 | 1 |
| | | | 0 |
| **3_prime_UTR_variant** | 53 | 73 | 126 |
| | | | 0 |
| splice_donor_variant | 2 | 10 | 12 |
| splice_acceptor_variant | 1 | 6 | 7 |
| splice_region_variant | 27 | 31 | 58 |
| | | | 0 |
| **intron_variant** | 179 | 210 | 389 |
| | | | 0 |
| NMD_transcript_variant | 121 | 227 | 348 |
| non_coding_transcript_exon_variant | 178 | 240 | 418 |
| non_coding_transcript_variant | 110 | 118 | 228 |
| Total (all Variant consequence entries) | 1028 | 1528 | 2556 |

**Supplementary Figure S1. Analysis of G4-Vars variant consequences. (A)** Diagram showing the location of each variant consequence term relative to the transcript structure calculated by Ensembl (based on the identification of all overlapping Ensembl transcripts and on the use of a rule-based approach to predict the effects that each allele of the variant may have on each transcript, using a set of consequence terms defined by the Sequence Ontology (SO)). The region of interest (proximal promoter region, PPR) is signalled. Adapted from https://m.ensembl.org/info/genome/variation/prediction/consequences.jpg. The region of interest of this work (proximal promoter region) is signalled. **(B)** Table with counts of variant consequence terms obtained in sub-groups A2 and B2 (from Supplementary Table S7). Notice that some G4-Vars ids can present different variant consequences in the same or in different genes.

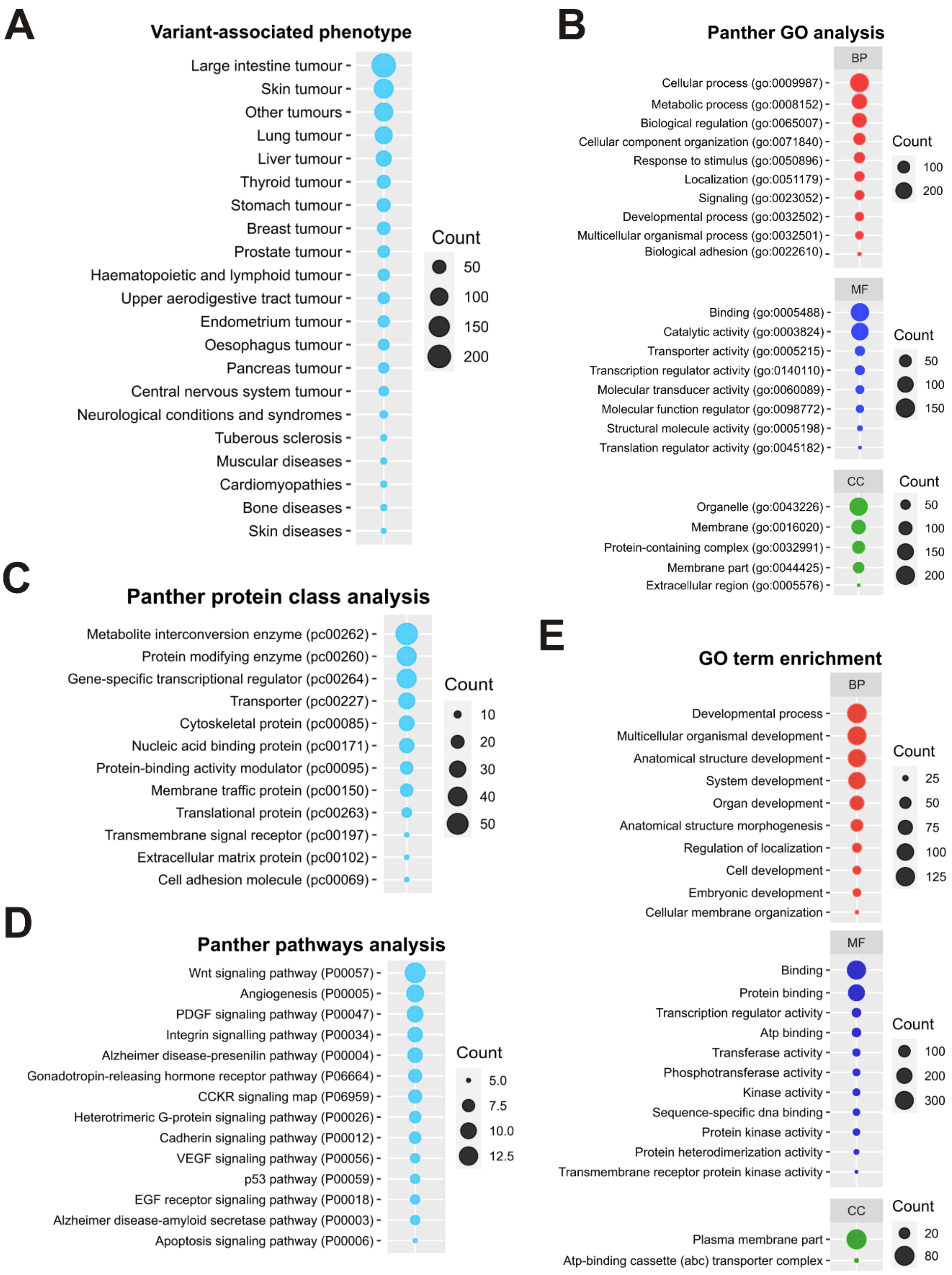


**Supplementary Figure S2.** ***In silico*** **characterization of G4-Vars and associated genes. (A)** Variant-associated phenotype characterization. We found 833 ids (out of 922) related to a variety of pathologies, the most represented being different kinds of tumours (intestinal, skin, lung, liver, stomach, and breast tumours, among others); although other conditions or diseases, such as neurological conditions and syndromes, cardiomyopathies, muscular, bone, and skin diseases were also found. The 587 genes associated with the 922 variants were characterised by considering

GO terms (biological process, BP; molecular function, MF; cellular component, CC) **(B)**, protein class **(C)**, and pathway **(D)**. **(E)** BiNGO gene ontology term enrichment study. Circles sizes represent the number of variants ids **(A)** or genes **(B-E)** related to each phenotype/GO term/protein class/pathway. Only statistically enriched GO terms are shown. Scales are shown on the right of each figure.

***In silico* characterization of G4-Vars and associated genes – Materials and Methods**

Variant ids corresponding to identified G4-Vars after Quadron predictions were used as inputs of the Biomart tool to download associated phenotype, genes, and gene features from Ensembl Variation and Ensembl Genes databases. G4-Vars-associated genes were characterised using Panther DB (http://pantherdb.org/) (3) regarding protein classes, biological pathways, and Gene Ontology (GO) terms. Besides, GO terms enrichment was analysed with the Biological Networks Gene Ontology (BiNGO) plugin using Cytoscape v3.2.1 (http://www.cytoscape.org/) (4). Statistical significance was tested using a hypergeometric test and a Benjamini & Hochberg false discovery rate correction (FDR). $p$-value $< 0.05$ was considered as significant after FDR correction.

### Supplementary Figure S3

# A

***GRIN2B***

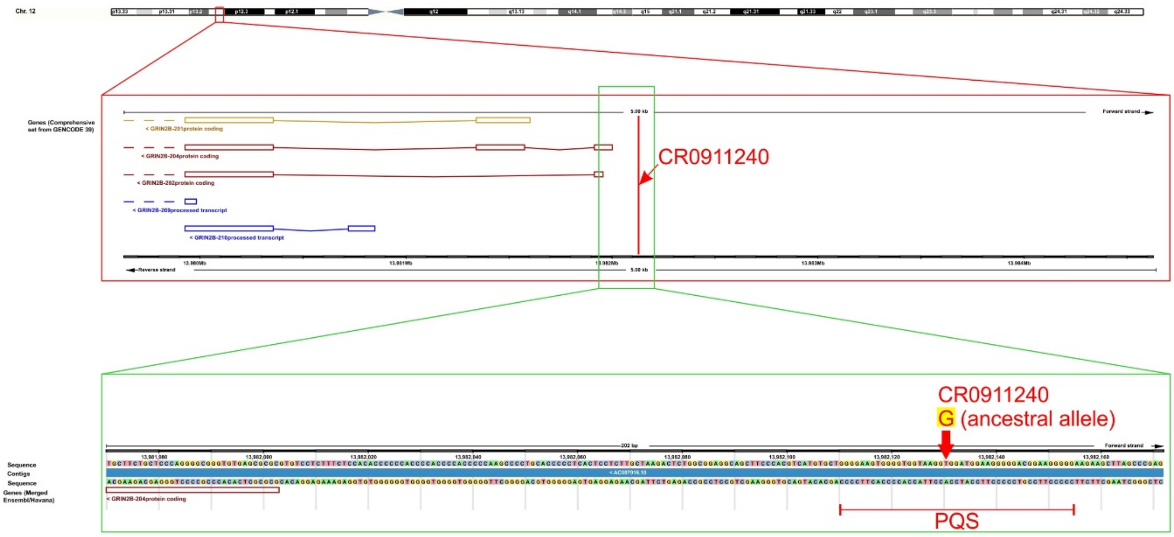


# B

***F7***

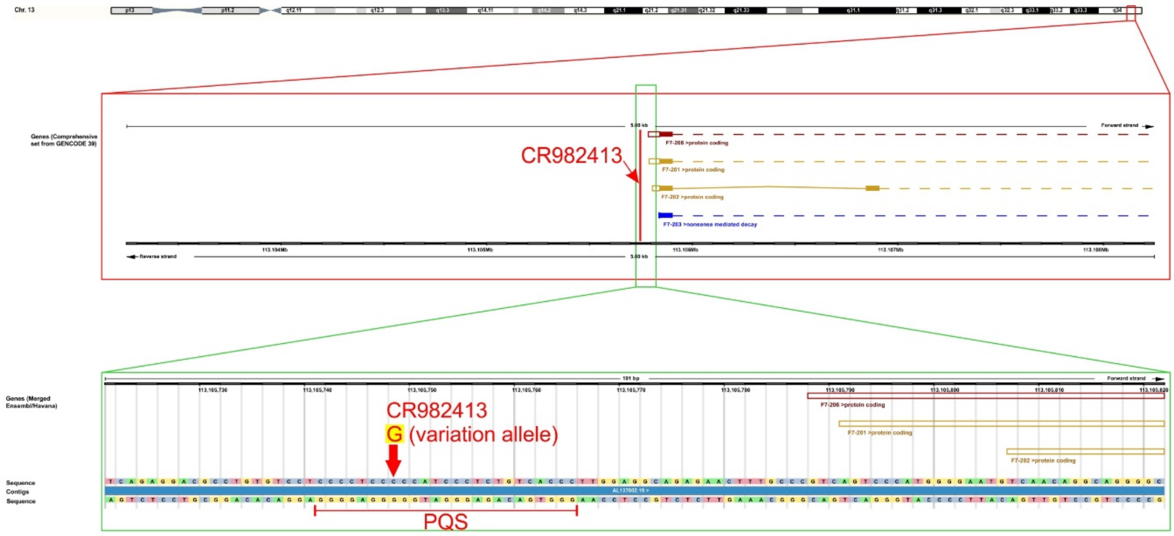

### Supplementary Figure S3 (cont.)

# C

#### *CSF2*

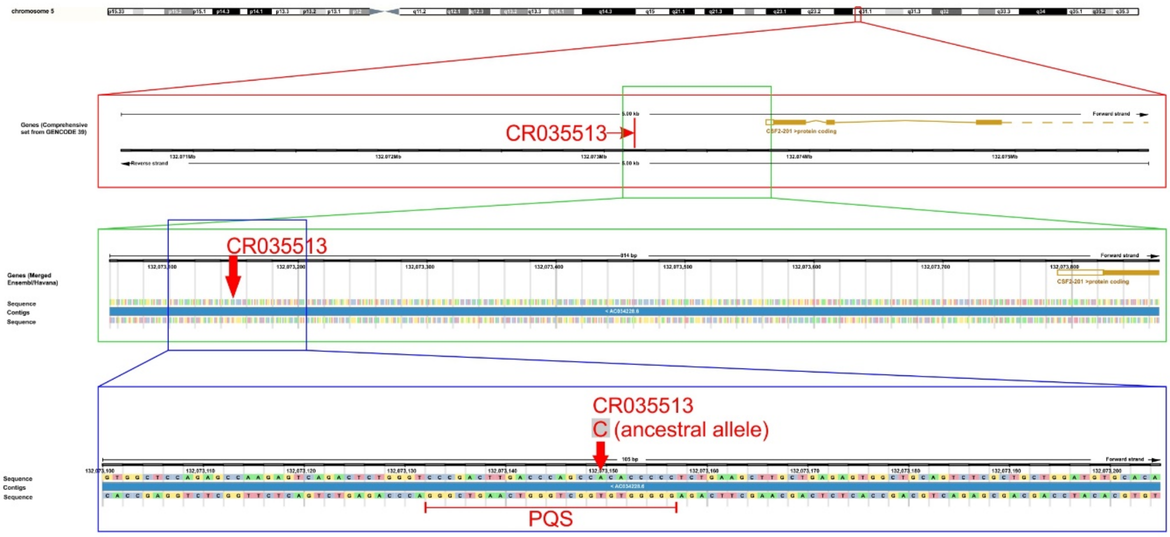


# D

#### *SIRT1*

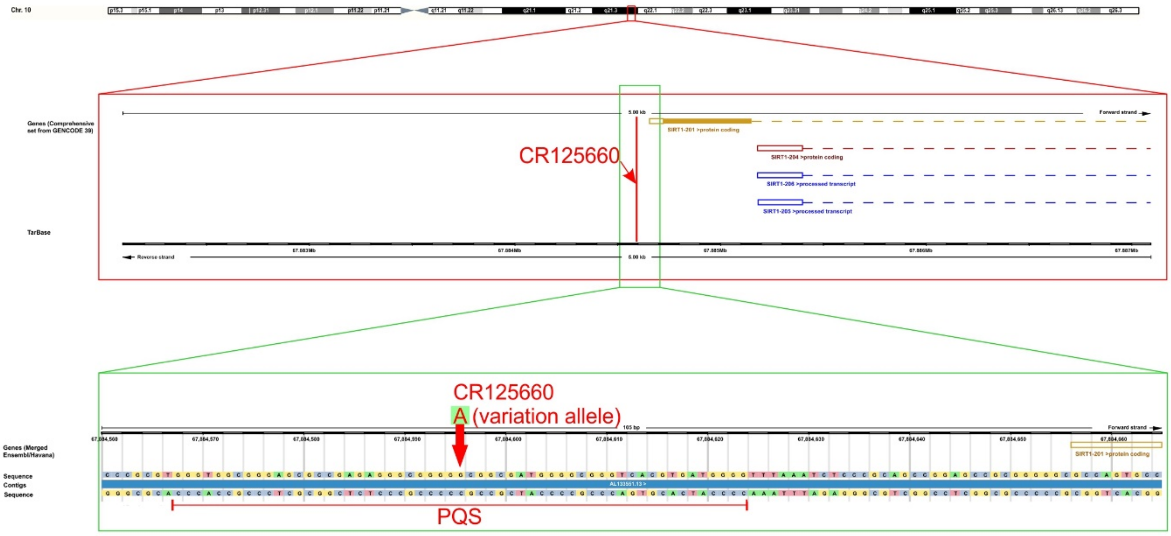

**Supplementary Figure S3 (cont.)**

**E**

***ZNF81***

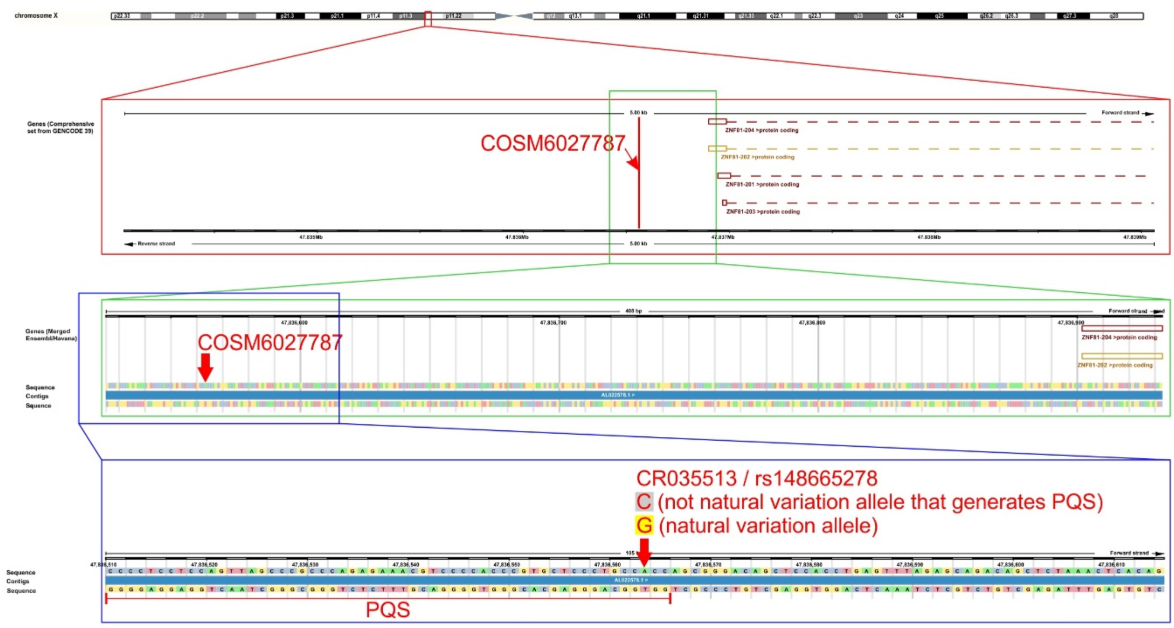


**F**

***LHFPL5***

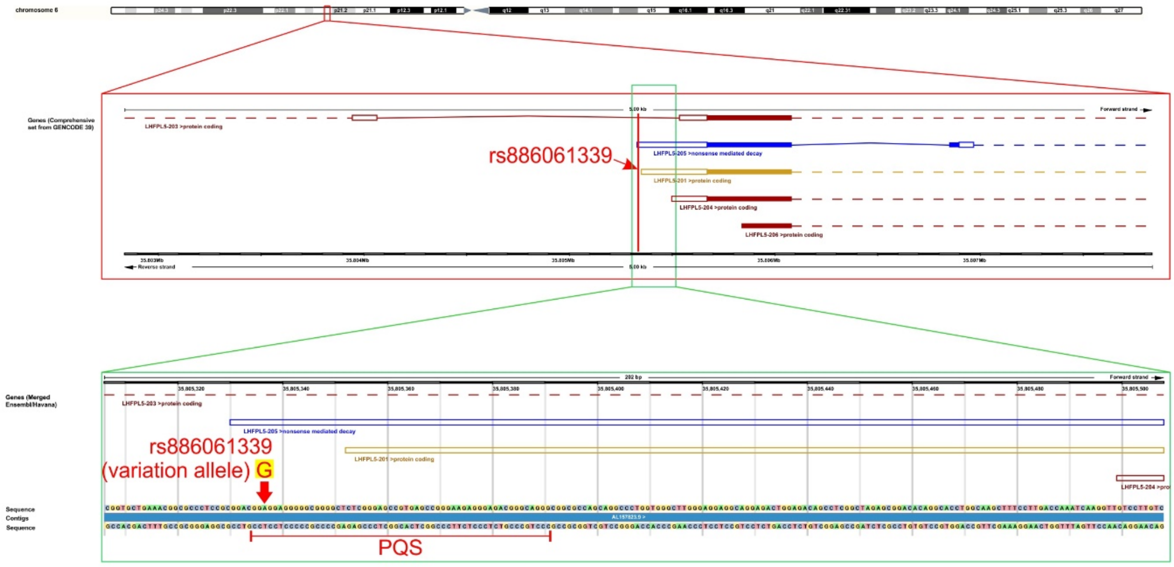


**Supplementary Figure S3. Details of the selected G4-Vars.** Ensembl screenshots of the selected G4-Vars from sub-groups A2 and B2 showing graphical schemes of chromosome position, genomic context with the nearby transcripts and detailed genomic region and nucleotide change. **(A)** CR0911240 for *GRIN2B*, **(B)** CR982413 for *F7*, **(C)** CR035513 for *CSF2*, **(D)** CR125660 for *SIRT1*, **(E)** COSM6027787* for *ZNF81* and **(F)** rs886061339** for *LHFPL5*.

* This G4-Var is classified in Ensembl as “upstream_gene_variant” and was retrieved from COSMIC catalogue, by replacing the nucleobase informed for the AA by the other

three possible nucleobases. Yet, this G4-Var was ruled out because the nucleobase replacement generating the PQS is not the one informed for a co-located variant in the dbSNP, thus suggesting it is not biologically relevant.

** This G4-Var is classified in Ensembl as "5_prime_UTR_variant" of a nonsense-mediated mRNA decay and is located in the first intron of an *LHFPL5* protein-coding transcript. It is also located upstream of the TSS (within the PPR) of other three *LHFPL5* protein-coding transcripts, including the one classified by Ensembl as the most biologically relevant transcript (LHFPL5-201).

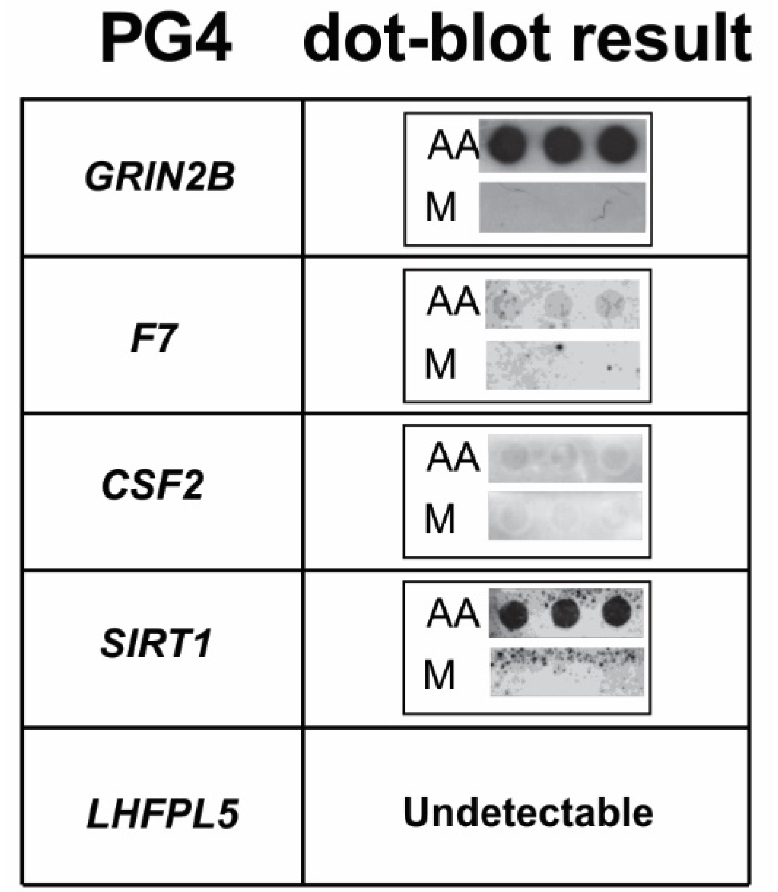


**Supplementary Figure S4. Immunodetection with BG4 antibody of G4s formation by PQSs of the selected G4-Vars.** AA- and mutated (M)-sequences containing the PQSs of each selected G4-Var were folded in 100 mM $K^+$ and dot-blotted on for immunodetection using BG4 antibody. VA- and M-sequences were blotted for the G4-Var of the sub-group B2 (*LHFPL5*).

**BG4 dot-blots – Materials and Methods**

Dots were spotted using 10 µl of 1 µM folded oligodeoxiribonucleotides (in 10 mM Tris pH 7.5 and 100 mM KCl) on Hybond™-N+ membrane (Amersham) and cross-linked with a UV lamp (UVP, 302 nm) during 5 min. Membranes were washed twice with Tris-Buffer Saline with 0.1% Tween 20 detergent (TBS-T) for 10 min at room temperature and then blocked in TBS-T containing 5% blotto for 1 h. Membranes were first incubated overnight at 4°C with 1 µg/ml of recombinant BG4-antibody (5). Then, membranes were incubated with Anti-Flag antibody (clone M2, Sigma Aldrich Catalogue Number F3165; 1 µg/ml) for 1 h at room temperature. Finally, membranes were incubated for 1 h at room temperature with anti-mouse conjugated to Horseradish Peroxidase (HRP) (Jackson ImmunoResearch Inc., Code: 115-035-003, dilution 1/25000). After each antibody incubation and before development, membranes were washed with TBS-T five times during 10 min at room temperature. Finally, G4s detection was developed by chemiluminescence (Biolumina Chemiluminescent Substrate, Kallium Technologies, CABA, Argentina) followed by exposition on X-ray films (Amersham Hyperfilm ECL, GE, MA, USA).

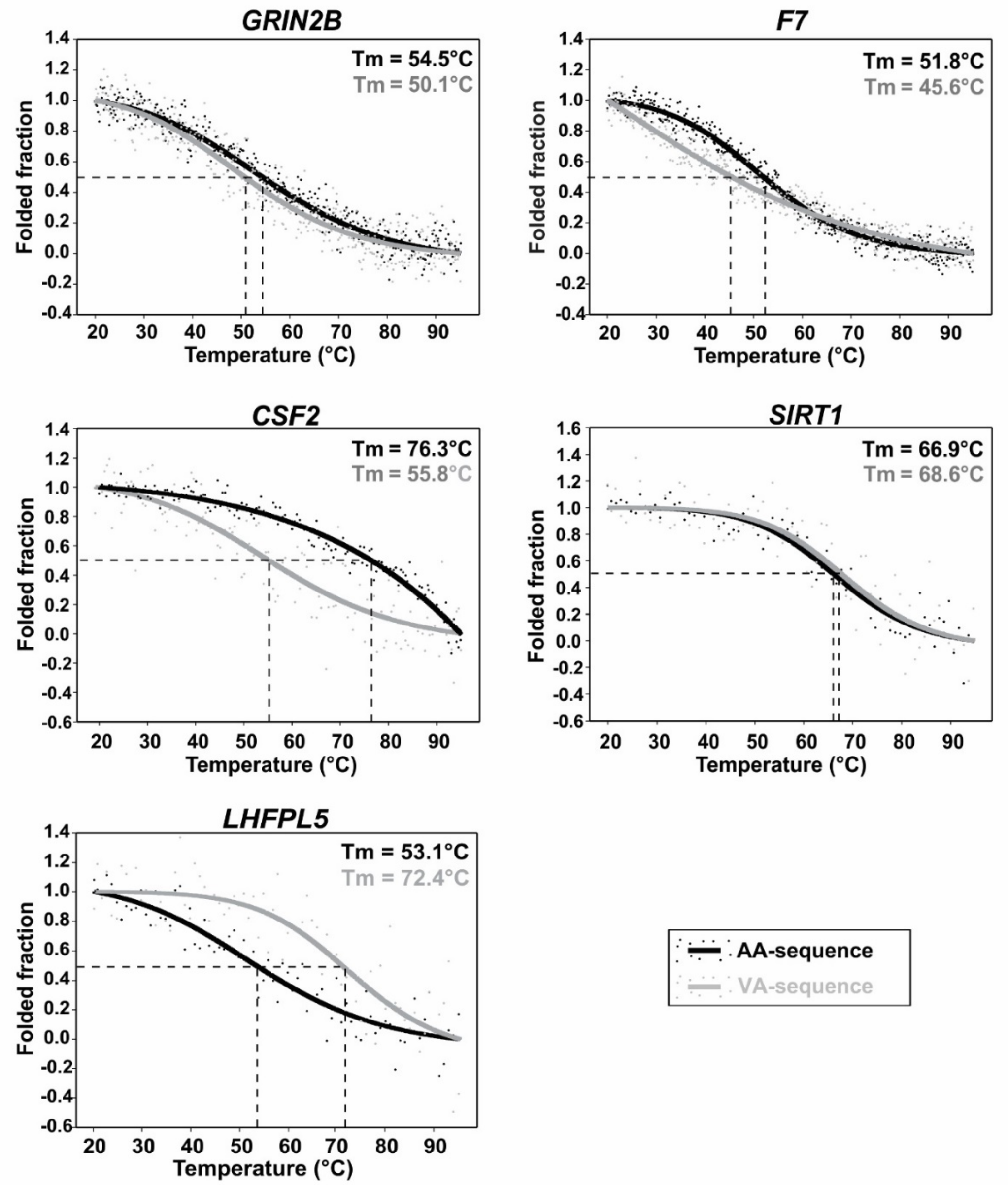


**Supplementary Figure S5. CD melting curves obtained for the PQSs of the selected G4-Vars.** Experimental data (dots) and fitted curves (solid lines) are represented. Estimated melting temperatures (Tm) are informed in the top-right corner of each plot box.

**CD melting curves – Materials and Methods**

Two µM oligodeoxyribonucleotides with the AA- and VA-sequences containing the PQSs of each selected G4-Var were folded in the presence of 100 mM KCl (as described for CD spectroscopy) prior to CD melting. CD melting curves were recorded by ellipticity measurements between 20°C and 95°C at the wavelength corresponding to the maximum observed in the spectra at the initial temperature (20°C) for the positive band around 264 nm, using the same parameters set for the spectra, a temperature increase speed of 1°C/min, and a sampling interval of 0.5°C. Data was analysed in SigmaPlot 11.0 as reported elsewhere (6) for calculation of melting temperatures (Tm).

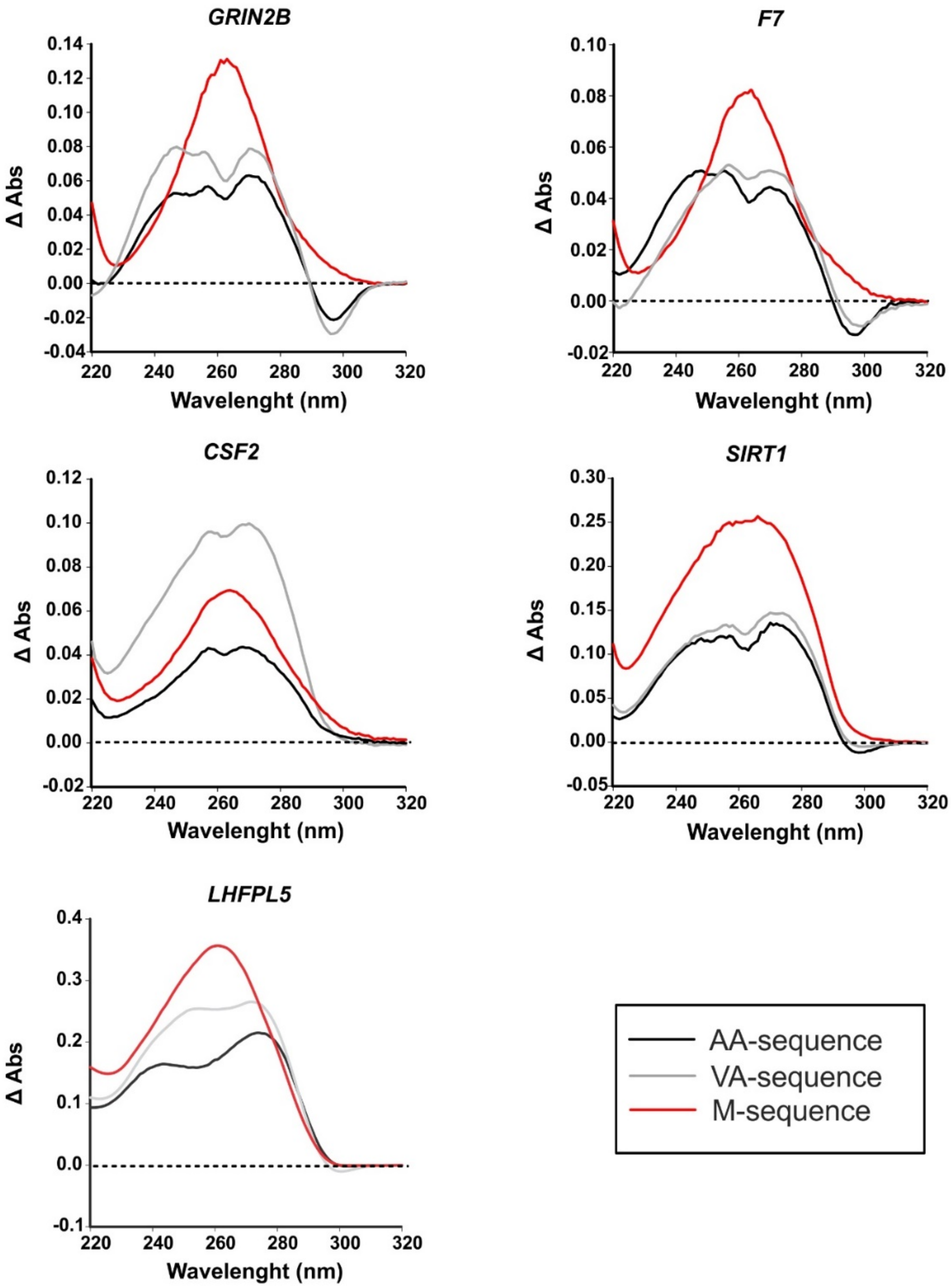


**Supplementary Figure S6. Thermal difference spectra (TDS) obtained for the PQSs of the selected G4-Vars.** TDS performed on AA-, VA- and M-sequences containing the PQSs of each selected G4-Var folded in the presence of 10 mM $K^+$ (*GRIN2B*), 100 mM $K^+$ (*F7*, *CSF2* and *SIRT1*) and 300 mM $K^+$ (*LHFPL5*) as defined from CD spectra (Figure 2).

**Thermal Difference Spectroscopy (TDS) – Materials and Methods**

TDS assays were performed as described elsewhere (6), with few modifications. Two μM oligodeoxyribonucleotides folded into G4 (as described for CD spectroscopy) were scanned to measure absorbance over the wavelength range of 220−320 nm (10 mm quartz cell, 1 nm band width, 1 nm data pitch, 100 nm/min scanning speed) at 40°C (folded condition) and then at 70°C (unfolded condition), using a Jasco V-630BIO spectrophotometer with peltier temperature control. The absorbance spectra obtained at 70°C and 40°C were subtracted (Δ Abs) and plotted on a graph to obtain TDS.

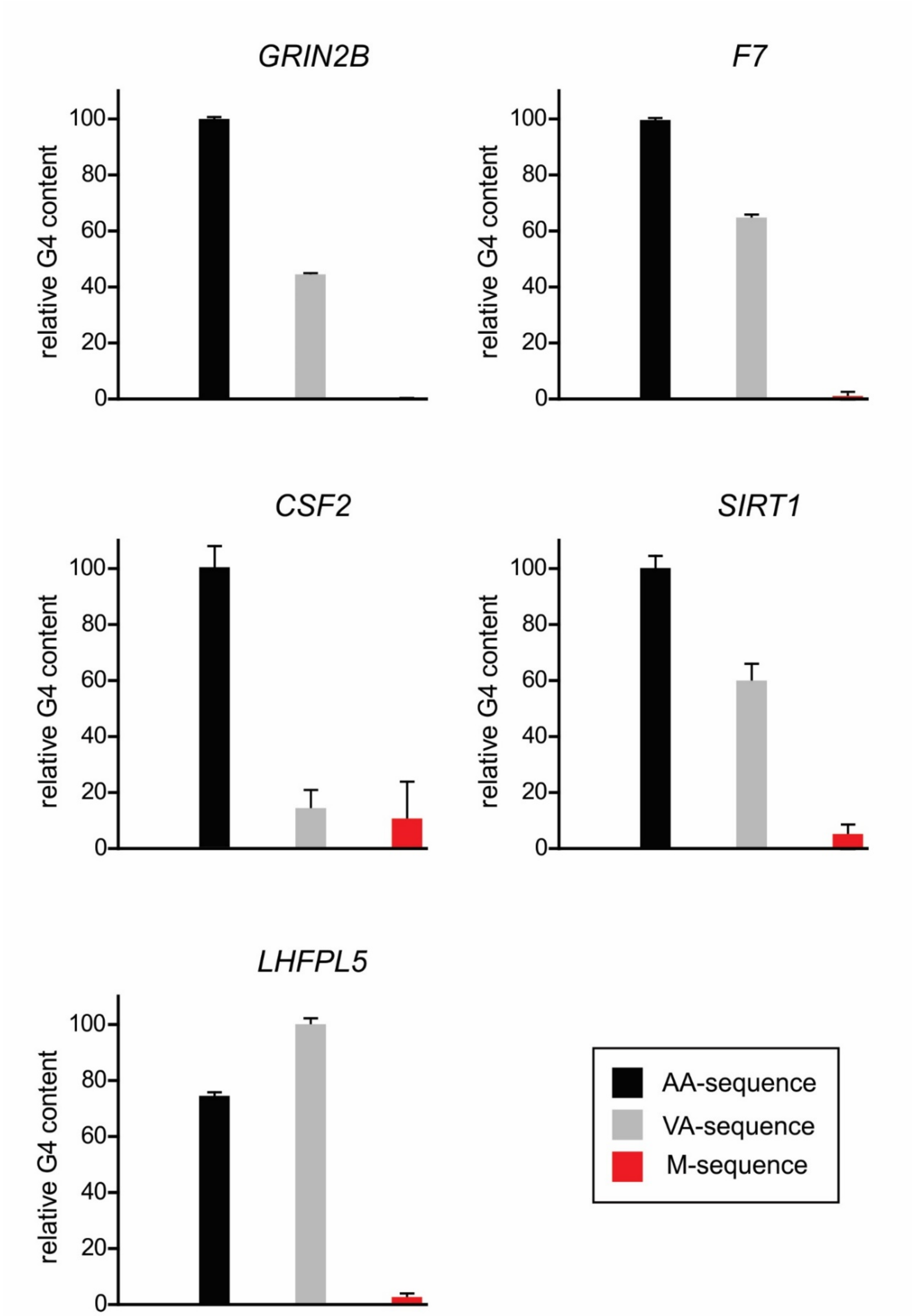


**Supplementary Figure S7. Relative G4 content determined by NMR.** Values were determined by NMR signal integration of the imino region in the 1D $^1$H NMR spectra depicted in Figures 3 and 4. The integral values for each oligodeoxyribonucleotide were referenced to the highest value within those for the same gene. Error bars represent the noise level for each determination.

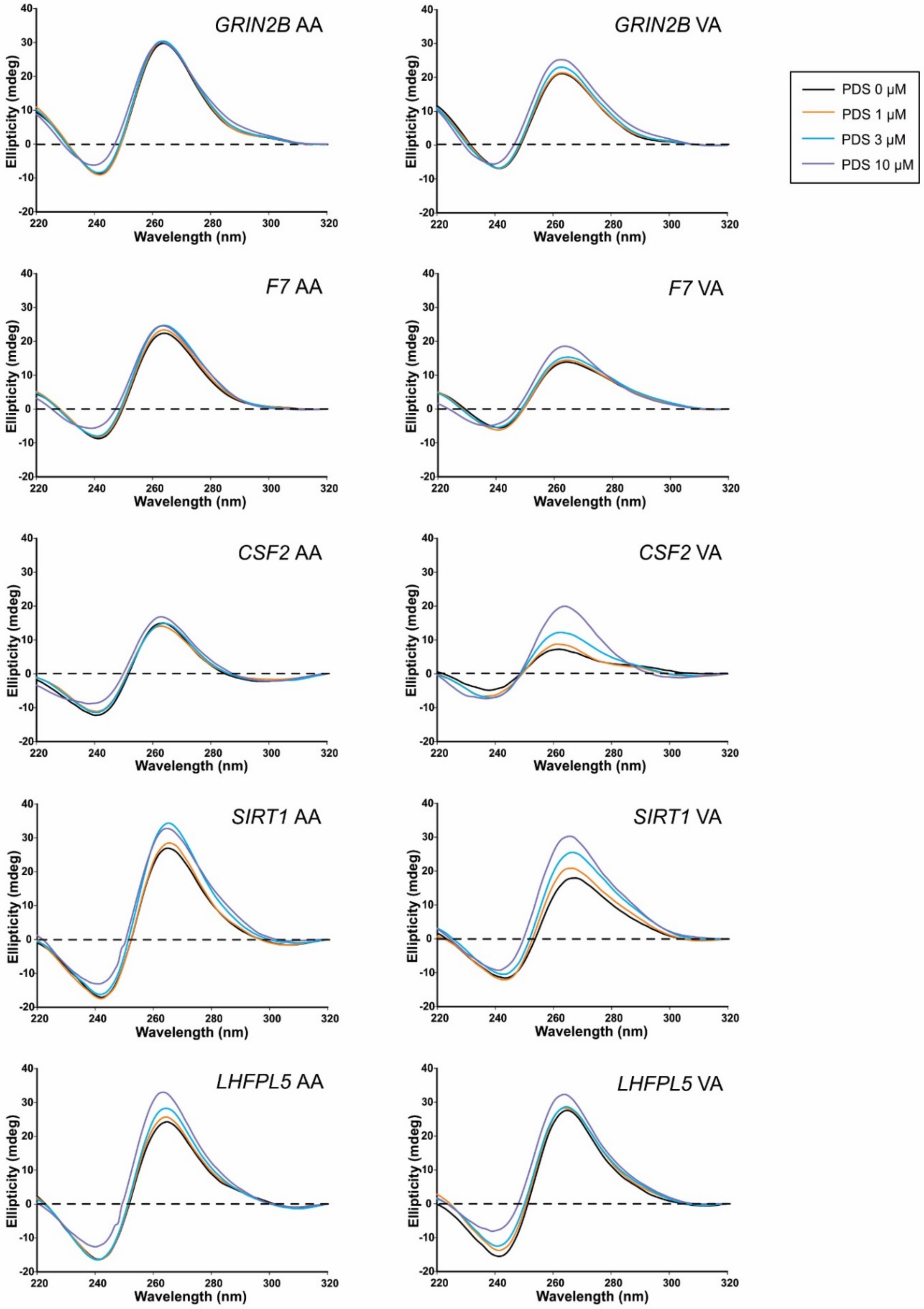


**Supplementary Figure S8. CD spectra of AA- and VA-sequences treated with PDS.** CD spectra performed for AA- and VA-sequences folded as in Figures 3 and 4 and then incubated in the absence or in the presence of 1, 3, and 10 µM pyridostatin (PDS, pyridostatin trifluoroacetate salt, Sigma-Aldrich SML0678) previous to recording the CD spectra.

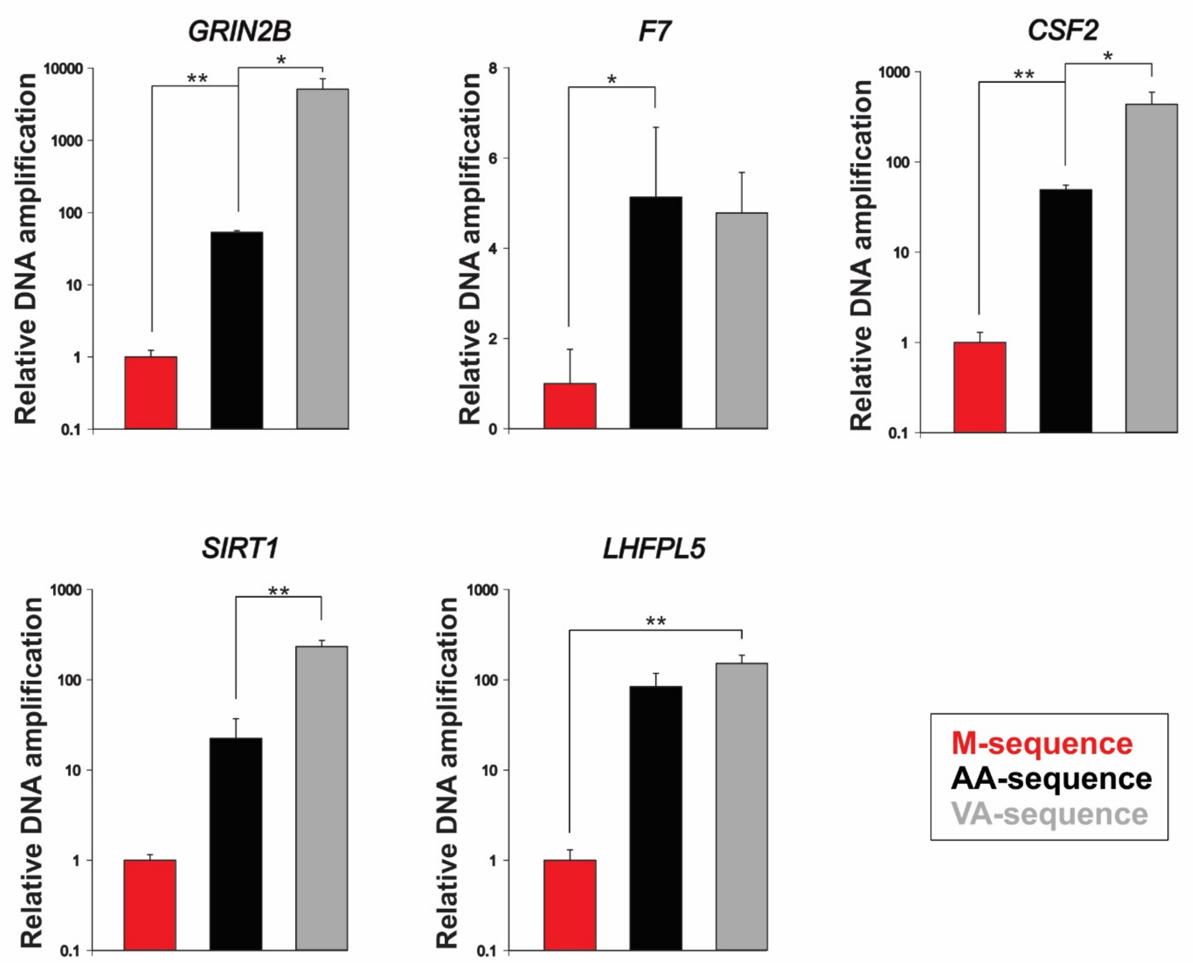


**Supplementary Figure S9. qPSA of AA-, VA- and M-sequences in the presence of PDS.** Bars represent the relative DNA amplification ($2^{-\Delta Cq}$) for AA-, VA-, and M-template sequences. ΔCq was defined as the threshold cycle in absence of PDS minus the threshold cycle in presence of 0.3 μM PDS (pyridostatin trifluoroacetate salt, Sigma-Aldrich SML0678) for the same sequence. Results were normalised to the relative DNA amplification of the corresponding M-sequence. $*p < 0,05$, $**p < 0.01$, T-Student test.

<u>**References for Supplementary Material**</u>
