## Supplementary File S5 for "Genetic variations in G-Quadruplex forming sequences affect the transcription of human disease-related genes"

***TFBS prediction in AA- and VA-sequences for the five selected G4-Vars***

#### PROMO software (1, 2) was used.

Default is 15% dissimilarity (85% similarity) -used in this work- but it can be modified by the user. The dissimilarity threshold is the parameter that controls how similar a sequence must be to the matrix to be reported as a hit.

Both strands of AA- and VA-sequences were searched for matches.

The nucleobases involved in the G4-Vars are signaled in red.

Whole data containing TF, TFBS, dissimilarity score and nucleobases involved in the G4-Vars which are differentially predicted between AA and VA-sequences are signaled in yellow.

**>GRIN2B AA-sequence**
GTGCTGGGGAAGTGGGGTGGTAAGGGGGATGGAAGGGGGACGGAAGGGGGAAGAA

-- Factors predicted by PROMO in this sequence ----------------------
NAME; MATRIX_WIDTH;
GR-alpha [T00337]; 5
STAT4 [T01577]; 6
c-Ets-1 [T00112]; 7
Elk-1 [T00250]; 9
TFII-I [T00824]; 6
MAZ [T00490]; 13
YY1 [T00915]; 4
PEA3 [T00685]; 9
RXR-alpha [T01345]; 7

-- PROMO predictions detail ------------------------------------------

Sequence name; Factor name; Start position; End position; Dissimilarity; String; RE equally; RE query
Sequence; GR-alpha [T00337]; 20; 24; 6.263098; TAAGG; 0.10742; 0.23507;
Sequence; GR-alpha [T00337]; 31; 35; 8.281568; GAAGG; 0.21484; 1.37075;
Sequence; GR-alpha [T00337]; 42; 46; 8.281568; GAAGG; 0.21484; 1.37075;
Sequence; STAT4 [T01577]; 7; 12; 2.941176; GGAAGT; 0.08057; 0.16736;
Sequence; STAT4 [T01577]; 30; 35; 5.882353; GGAAGG; 0.01343; 0.23092;
Sequence; STAT4 [T01577]; 41; 46; 5.882353; GGAAGG; 0.01343; 0.23092;
Sequence; STAT4 [T01577]; 48; 53; 4.411765; GGAAGA; 0.05371; 0.16797;
Sequence; c-Ets-1 [T00112]; 5; 11; 5.558311; GGGGAAG; 0.01007; 0.20573;
Sequence; c-Ets-1 [T00112]; 28; 34; 4.910652; ATGGAAG; 0.01343; 0.04112;
Sequence; c-Ets-1 [T00112]; 39; 45; 3.359159; ACGGAAG; 0.01678; 0.01040;
Sequence; c-Ets-1 [T00112]; 46; 52; 5.558311; GGGGAAG; 0.01007; 0.20573;
Sequence; Elk-1 [T00250]; 3; 11; 12.179139; CTGGGGAAG; 0.00671; 0.03916;
Sequence; Elk-1 [T00250]; 26; 34; 9.620020; GGATGGAAG; 0.00210; 0.02605;
Sequence; Elk-1 [T00250]; 37; 45; 7.420684; GGACGGAAG; 0.00084; 0.00214;
Sequence; Elk-1 [T00250]; 44; 52; 8.931691; AGGGGGAAG; 0.00671; 0.10680;
Sequence; TFII-I [T00824]; 7; 12; 14.269340; GGAAGT; 0.16113; 0.10328;
Sequence; TFII-I [T00824]; 26; 31; 9.512894; GGATGG; 0.20142; 0.47214;
Sequence; TFII-I [T00824]; 30; 35; 9.512894; GGAAGG; 0.20142; 0.47214;
Sequence; TFII-I [T00824]; 37; 42; 9.512894; GGACGG; 0.20142; 0.47214;
Sequence; TFII-I [T00824]; 41; 46; 9.512894; GGAAGG; 0.20142; 0.47214;
Sequence; MAZ [T00490]; 21; 33; 14.932273; AAGGGGGATGGAA; 0.00176; 0.37110;
Sequence; MAZ [T00490]; 32; 44; 14.932273; AAGGGGGACGGAA; 0.00176; 0.37110;
Sequence; YY1 [T00915]; 28; 31; 0.000000; ATGG; 0.21484; 0.27592;
Sequence; PEA3 [T00685]; 25; 33; 12.454190; GGGATGGAA; 0.00084; 0.00433;
Sequence; RXR-alpha [T01345]; 14; 20; 6.967687; GGGTGGT; 0.01007; 0.02721;

**>GRIN2B VA-sequence**
GTGCTGGGGAAGTGGGGTGGTAAGGTGGATGGAAGGGGGACGGAAGGGGGAAGAA

-- Factors predicted by PROMO in this sequence ----------------------
NAME; MATRIX_WIDTH;
GR-alpha [T00337]; 5
STAT4 [T01577]; 6
c-Ets-1 [T00112]; 7
Elk-1 [T00250]; 9
TFII-I [T00824]; 6
MAZ [T00490]; 13
YY1 [T00915]; 4
PEA3 [T00685]; 9
RXR-alpha [T01345]; 7

-- PROMO predictions detail ------------------------------------------

Sequence name; Factor name; Start position; End position; Dissimilarity; String; RE equally; RE query
Sequence; GR-alpha [T00337]; 20; 24; 6.263098; TAAGG; 0.10742; 0.25243;
Sequence; GR-alpha [T00337]; 31; 35; 8.281568; GAAGG; 0.21484; 1.22560;
Sequence; GR-alpha [T00337]; 42; 46; 8.281568; GAAGG; 0.21484; 1.22560;
Sequence; STAT4 [T01577]; 7; 12; 2.941176; GGAAGT; 0.08057; 0.16306;
Sequence; STAT4 [T01577]; 30; 35; 5.882353; GGAAGG; 0.01343; 0.20418;
Sequence; STAT4 [T01577]; 41; 46; 5.882353; GGAAGG; 0.01343; 0.20418;
Sequence; STAT4 [T01577]; 48; 53; 4.411765; GGAAGA; 0.05371; 0.15955;
Sequence; c-Ets-1 [T00112]; 5; 11; 5.558311; GGGGAAG; 0.01007; 0.17819;
Sequence; c-Ets-1 [T00112]; 28; 34; 4.910652; ATGGAAG; 0.01343; 0.04376;
Sequence; c-Ets-1 [T00112]; 39; 45; 3.359159; ACGGAAG; 0.01678; 0.00978;
Sequence; c-Ets-1 [T00112]; 46; 52; 5.558311; GGGGAAG; 0.01007; 0.17819;
Sequence; Elk-1 [T00250]; 3; 11; 12.179139; CTGGGGAAG; 0.00671; 0.03801;
Sequence; Elk-1 [T00250]; 26; 34; 9.620020; GGATGGAAG; 0.00210; 0.02239;
Sequence; Elk-1 [T00250]; 37; 45; 7.420684; GGACGGAAG; 0.00084; 0.00189;
Sequence; Elk-1 [T00250]; 44; 52; 8.931691; AGGGGGAAG; 0.00671; 0.09180;
Sequence; TFII-I [T00824]; 7; 12; 14.269340; GGAAGT; 0.16113; 0.11616;
Sequence; TFII-I [T00824]; 26; 31; 9.512894; GGATGG; 0.20142; 0.44662;
Sequence; TFII-I [T00824]; 30; 35; 9.512894; GGAAGG; 0.20142; 0.44662;
Sequence; TFII-I [T00824]; 37; 42; 9.512894; GGACGG; 0.20142; 0.44662;
Sequence; TFII-I [T00824]; 41; 46; 9.512894; GGAAGG; 0.20142; 0.44662;
Sequence; MAZ [T00490]; 32; 44; 14.932273; AAGGGGGACGGAA; 0.00176; 0.30435;
Sequence; YY1 [T00915]; 28; 31; 0.000000; ATGG; 0.21484; 0.30276;
Sequence; PEA3 [T00685]; 25; 33; 9.340643; TGGATGGAA; 0.00084; 0.00268;

**>F7 AA-sequence**

TCCAAGGGTGACAGAGGGATGGGGGAGGGGAGGAC

-- Factors predicted by PROMO in this sequence ----------------------
NAME; MATRIX_WIDTH;
GR-alpha [T00337]; 5
NFI/CTF [T00094]; 8
TFII-I [T00824]; 6
MAZ [T00490]; 13
YY1 [T00915]; 4
PEA3 [T00685]; 9
c-Jun [T00133]; 7
RXR-alpha [T01345]; 7
T3R-beta1 [T00851]; 9
C/EBPbeta [T00581]; 4

-- PROMO predictions detail ------------------------------------------

Sequence name; Factor name; Start position; End position; Dissimilarity; String; RE equally; RE query
Sequence; GR-alpha [T00337]; 2; 6; 8.281568; CAAGG; 0.13672; 0.69974;
Sequence; GR-alpha [T00337]; 12; 16; 0.207689; AGAGG; 0.13672; 0.34268;
Sequence; GR-alpha [T00337]; 23; 27; 8.281568; GGAGG; 0.13672; 0.69974;
Sequence; GR-alpha [T00337]; 28; 32; 8.281568; GGAGG; 0.13672; 0.69974;
Sequence; NFI/CTF [T00094]; 1; 8; 3.793671; CCAAGGGT; 0.00320; 0.00097;
Sequence; TFII-I [T00824]; 16; 21; 9.512894; GGATGG; 0.12817; 0.28200;
Sequence; TFII-I [T00824]; 23; 28; 11.337888; GGAGGG; 0.04272; 0.33129;
Sequence; MAZ [T00490]; 18; 30; 3.986869; ATGGGGGAGGGGA; 0.00010; 0.00838;
Sequence; YY1 [T00915]; 18; 21; 0.000000; ATGG; 0.13672; 0.11871;
Sequence; PEA3 [T00685]; 15; 23; 13.648823; GGGATGGGG; 0.00481; 0.01491;
Sequence; c-Jun [T00133]; 8; 14; 6.787369; TGACAGA; 0.01282; 0.00466;
Sequence; RXR-alpha [T01345]; 5; 11; 5.937582; GGGTGAC; 0.01282; 0.02797;
Sequence; T3R-beta1 [T00851]; 2; 10; 7.774776; CAAGGGTGA; 0.00481; 0.01115;
Sequence; C/EBPbeta [T00581]; 1; 4; 1.639871; CCAA; 0.27344; 0.08229;

**>F7 VA-sequence**
TCCAAGGGTGACAGAGGGATGGCGGAGGGGAGGAC

-- Factors predicted by PROMO in this sequence ----------------------
NAME; MATRIX_WIDTH;
GR-alpha [T00337]; 5
NFI/CTF [T00094]; 8
XBP-1 [T00902]; 6
ENKTF-1 [T00255]; 8
TFII-I [T00824]; 6
E2F-1 [T01542]; 8
YY1 [T00915]; 4
PEA3 [T00685]; 9
c-Jun [T00133]; 7
RXR-alpha [T01345]; 7
T3R-beta1 [T00851]; 9
C/EBPbeta [T00581]; 4

-- PROMO predictions detail ------------------------------------------

Sequence name; Factor name; Start position; End position; Dissimilarity; String; RE equally; RE query
Sequence; GR-alpha [T00337]; 2; 6; 8.281568; CAAGG; 0.13672; 0.60796;
Sequence; GR-alpha [T00337]; 12; 16; 0.207689; AGAGG; 0.13672; 0.30868;
Sequence; GR-alpha [T00337]; 23; 27; 8.281568; GGAGG; 0.13672; 0.60796;
Sequence; GR-alpha [T00337]; 28; 32; 8.281568; GGAGG; 0.13672; 0.60796;
Sequence; NFI/CTF [T00094]; 1; 8; 3.793671; CCAAGGGT; 0.00320; 0.00148;
Sequence; XBP-1 [T00902]; 18; 23; 9.789909; ATGGCG; 0.03418; 0.01474;
Sequence; ENKTF-1 [T00255]; 19; 26; 1.255756; TGGCGGAG; 0.00427; 0.00678;
Sequence; TFII-I [T00824]; 16; 21; 9.512894; GGATGG; 0.12817; 0.26463;
Sequence; TFII-I [T00824]; 23; 28; 11.337888; GGAGGG; 0.04272; 0.27525;
Sequence; E2F-1 [T01542]; 21; 28; 12.813402; GCGGAGGG; 0.00801; 0.03405;
Sequence; YY1 [T00915]; 18; 21; 0.000000; ATGG; 0.13672; 0.10989;
Sequence; PEA3 [T00685]; 15; 23; 13.648823; GGGATGGCG; 0.00481; 0.01291;
Sequence; c-Jun [T00133]; 8; 14; 6.787369; TGACAGA; 0.01282; 0.00564;
Sequence; RXR-alpha [T01345]; 5; 11; 5.937582; GGGTGAC; 0.01282; 0.02508;
Sequence; T3R-beta1 [T00851]; 2; 10; 7.774776; CAAGGGTGA; 0.00481; 0.00951;
Sequence; C/EBPbeta [T00581]; 1; 4; 1.639871; CCAA; 0.27344; 0.08879;

**>CSF2 AA-sequence**

TCAGAGGGGGTGGGGCTGGGTCAAGTCGGGACCCA

-- Factors predicted by PROMO in this sequence ----------------------
NAME; MATRIX_WIDTH;
GR-alpha [T00337]; 5
c-Jun [T00133]; 7
RXR-alpha [T01345]; 7
C/EBPbeta [T00581]; 4
ER-alpha [T00261]; 5
RAR-alpha1 [T00719]; 13
PPAR-alpha:RXR-alpha [T05221]; 11
Pax-5 [T00070]; 7
p53 [T00671]; 7
ETF [T00270]; 11

-- PROMO predictions detail ------------------------------------------

Sequence name; Factor name; Start position; End position; Dissimilarity; String; RE equally; RE query
Sequence; GR-alpha [T00337]; 2; 6; 0.207689; AGAGG; 0.13672; 0.13562;
Sequence; c-Jun [T00133]; 16; 22; 3.049104; TGGGTCA; 0.00427; 0.00245;
Sequence; RXR-alpha [T01345]; 7; 13; 5.271235; GGGTGGG; 0.01068; 0.04894;
Sequence; RXR-alpha [T01345]; 17; 23; 0.848226; GGGTCAA; 0.00854; 0.01260;
Sequence; RXR-alpha [T01345]; 27; 33; 4.241130; GGGACCC; 0.01709; 0.03124;
Sequence; C/EBPbeta [T00581]; 20; 23; 1.366559; TCAA; 0.27344; 0.14926;
Sequence; ER-alpha [T00261]; 18; 22; 0.000000; GGTCA; 0.03418; 0.02855;
Sequence; RAR-alpha1 [T00719]; 16; 28; 13.441672; TGGGTCAAGTCGG; 0.00055; 0.00097;
Sequence; PPAR-alpha:RXR-alpha [T05221]; 13; 23; 4.727619; GGCTGGGTCAA; 0.00057; 0.00090;
Sequence; Pax-5 [T00070]; 12; 18; 0.000000; GGGCTGG; 0.01923; 0.09233;
Sequence; p53 [T00671]; 12; 18; 3.750231; GGGCTGG; 0.01282; 0.05545;
Sequence; ETF [T00270]; 5; 15; 7.870358; GGGGGTGGGGC; 0.00125; 0.02659;

**>CSF2 VA-sequence**

TCAGAGGGGGTGTGGCTGGGTCAAGTCGGGACCCA

-- Factors predicted by PROMO in this sequence ----------------------
NAME; MATRIX_WIDTH;
GR-alpha [T00337]; 5
ENKTF-1 [T00255]; 8
c-Jun [T00133]; 7
RXR-alpha [T01345]; 7
C/EBPbeta [T00581]; 4
ER-alpha [T00261]; 5
RAR-alpha1 [T00719]; 13
PPAR-alpha:RXR-alpha [T05221]; 11

-- PROMO predictions detail ------------------------------------------

Sequence name; Factor name; Start position; End position; Dissimilarity; String; RE equally; RE query
Sequence; GR-alpha [T00337]; 2; 6; 0.207689; AGAGG; 0.13672; 0.12805;
Sequence; ENKTF-1 [T00255]; 12; 19; 8.198520; TGGCTGGG; 0.01282; 0.02999;
Sequence; c-Jun [T00133]; 16; 22; 3.049104; TGGGTCA; 0.00427; 0.00291;
Sequence; RXR-alpha [T01345]; 7; 13; 4.019014; GGGTGTG; 0.01709; 0.03228;
Sequence; RXR-alpha [T01345]; 17; 23; 0.848226; GGGTCAA; 0.00854; 0.01369;
Sequence; RXR-alpha [T01345]; 27; 33; 4.241130; GGGACCC; 0.01709; 0.03228;
Sequence; C/EBPbeta [T00581]; 20; 23; 1.366559; TCAA; 0.27344; 0.18598;
Sequence; ER-alpha [T00261]; 18; 22; 0.000000; GGTCA; 0.03418; 0.03090;
Sequence; RAR-alpha1 [T00719]; 16; 28; 13.441672; TGGGTCAAGTCGG; 0.00055; 0.00092;
Sequence; PPAR-alpha:RXR-alpha [T05221]; 13; 23; 4.727619; GGCTGGGTCAA; 0.00057; 0.00090;

**>SIRT1 AA-sequence**

CGCGTGGGTGGCGGGAGCGCCGAGAGGGCGGGGGCGGCGATGGGGCGGGTCACGTGATGGGGTTTAA

-- Factors predicted by PROMO in this sequence ----------------------
NAME; MATRIX_WIDTH;
ENKTF-1 [T00255]; 8
GR-alpha [T00337]; 5
c-Jun [T00133]; 7
c-Myc [T00140]; 6
ATF-1 [T00968]; 11
USF1 [T00874]; 10
FOXP3 [T04280]; 6
RXR-alpha [T01345]; 7
YY1 [T00915]; 4
ER-alpha [T00261]; 5
RAR-alpha1 [T00719]; 13
E2F-1 [T01542]; 8
Pax-5 [T00070]; 7
p53 [T00671]; 7
Sp1 [T00759]; 10
MAZ [T00490]; 13
WT1 [T00899]; 9
GCF [T00320]; 9
TFII-I [T00824]; 6

-- PROMO predictions detail ------------------------------------------

Sequence name; Factor name; Start position; End position; Dissimilarity; String; RE equally; RE query
Sequence; ENKTF-1 [T00255]; 8; 15; 8.198520; TGGCGGGA; 0.02454; 0.07022;
Sequence; GR-alpha [T00337]; 22; 26; 0.207689; AGAGG; 0.26172; 0.17303;
Sequence; c-Jun [T00133]; 45; 51; 7.178905; CGGGTCA; 0.02454; 0.00941;
Sequence; c-Myc [T00140]; 50; 55; 0.000000; CACGTG; 0.01636; 0.01109;
Sequence; ATF-1 [T00968]; 46; 56; 5.246906; GGGTCACGTGA; 0.00096; 0.00039;
Sequence; USF1 [T00874]; 46; 55; 3.987093; GGGTCACGTG; 0.00268; 0.00180;
Sequence; USF1 [T00874]; 50; 59; 4.677985; CACGTGATGG; 0.00230; 0.00054;
Sequence; FOXP3 [T04280]; 61; 66; 11.337888; GTTTAA; 0.08179; 0.02231;
Sequence; RXR-alpha [T01345]; 5; 11; 7.815913; GGGTGGC; 0.00818; 0.05525;
Sequence; RXR-alpha [T01345]; 46; 52; 4.241130; GGGTCAC; 0.03271; 0.07289;
Sequence; RXR-alpha [T01345]; 59; 65; 0.626110; GGGTTTA; 0.00409; 0.00181;
Sequence; YY1 [T00915]; 39; 42; 0.000000; ATGG; 0.26172; 0.19008;
Sequence; YY1 [T00915]; 56; 59; 0.000000; ATGG; 0.26172; 0.19008;
Sequence; ER-alpha [T00261]; 47; 51; 0.000000; GGTCA; 0.06543; 0.04073;
Sequence; RAR-alpha1 [T00719]; 45; 57; 12.025302; CGGGTCACGTGAT; 0.00008; 0.00003;
Sequence; E2F-1 [T01542]; 10; 17; 5.042045; GCGGGAGC; 0.00613; 0.04647;
Sequence; E2F-1 [T01542]; 27; 34; 10.518902; GCGGGGGC; 0.02045; 0.25113;
Sequence; E2F-1 [T01542]; 33; 40; 11.323028; GCGGCGAT; 0.01636; 0.06258;
Sequence; E2F-1 [T01542]; 44; 51; 7.839654; GCGGGTCA; 0.01022; 0.01937;
Sequence; Pax-5 [T00070]; 25; 31; 1.537547; GGGCGGG; 0.02454; 0.34735;
Sequence; Pax-5 [T00070]; 31; 37; 9.552105; GGGCGGC; 0.04907; 0.20040;
Sequence; Pax-5 [T00070]; 42; 48; 1.537547; GGGCGGG; 0.02454; 0.34735;
Sequence; p53 [T00671]; 25; 31; 3.375208; GGGCGGG; 0.02454; 0.30574;
Sequence; p53 [T00671]; 31; 37; 6.188498; GGGCGGC; 0.02045; 0.10413;
Sequence; p53 [T00671]; 42; 48; 3.375208; GGGCGGG; 0.02454; 0.30574;
Sequence; Sp1 [T00759]; 24; 33; 1.150552; AGGGCGGGGG; 0.00045; 0.03319;
Sequence; Sp1 [T00759]; 30; 39; 2.729105; GGGGCGGCGA; 0.00134; 0.04644;
Sequence; Sp1 [T00759]; 41; 50; 1.229271; GGGGCGGGTC; 0.00045; 0.03319;
Sequence; MAZ [T00490]; 38; 50; 14.398434; GATGGGGCGGGTC; 0.00034; 0.03163;
Sequence; WT1 [T00899]; 10; 18; 11.111111; GCGGGAGCG; 0.00690; 0.29648;
Sequence; WT1 [T00899]; 27; 35; 0.000000; GCGGGGGCG; 0.00026; 0.02035;
Sequence; GCF [T00320]; 16; 24; 4.846987; GCGCCGAGA; 0.00920; 0.03968;
Sequence; TFII-I [T00824]; 13; 18; 11.337888; GGAGCG; 0.08179; 0.35432;

**>SIRT1 VA-sequence**

CGCGTGGGTGGCGGGAGCGCCGAGAGGGCGGGGACGGCGATGGGGCGGGTCACGTGATGGGGTTTAA

-- Factors predicted by PROMO in this sequence ----------------------
NAME; MATRIX_WIDTH;
ENKTF-1 [T00255]; 8
GR-alpha [T00337]; 5
c-Jun [T00133]; 7
c-Myc [T00140]; 6
ATF-1 [T00968]; 11
USF1 [T00874]; 10
FOXP3 [T04280]; 6
RXR-alpha [T01345]; 7
YY1 [T00915]; 4
ER-alpha [T00261]; 5
RAR-alpha1 [T00719]; 13
E2F-1 [T01542]; 8
Pax-5 [T00070]; 7
p53 [T00671]; 7
Sp1 [T00759]; 10
MAZ [T00490]; 13
TFII-I [T00824]; 6
WT1 [T00899]; 9
GCF [T00320]; 9

-- PROMO predictions detail ------------------------------------------

Sequence name; Factor name; Start position; End position; Dissimilarity; String; RE equally; RE query
Sequence; ENKTF-1 [T00255]; 8; 15; 8.198520; TGGCGGGA; 0.02454; 0.06641;
Sequence; GR-alpha [T00337]; 22; 26; 0.207689; AGAGG; 0.26172; 0.20374;
Sequence; c-Jun [T00133]; 45; 51; 7.178905; CGGGTCA; 0.02454; 0.01049;
Sequence; c-Myc [T00140]; 50; 55; 0.000000; CACGTG; 0.01636; 0.01183;
Sequence; ATF-1 [T00968]; 46; 56; 5.246906; GGGTCACGTGA; 0.00096; 0.00044;
Sequence; USF1 [T00874]; 46; 55; 3.987093; GGGTCACGTG; 0.00268; 0.00196;
Sequence; USF1 [T00874]; 50; 59; 4.677985; CACGTGATGG; 0.00230; 0.00063;
Sequence; FOXP3 [T04280]; 61; 66; 11.337888; GTTTAA; 0.08179; 0.02432;
Sequence; RXR-alpha [T01345]; 5; 11; 7.815913; GGGTGGC; 0.00818; 0.04949;
Sequence; RXR-alpha [T01345]; 46; 52; 4.241130; GGGTCAC; 0.03271; 0.06955;
Sequence; RXR-alpha [T01345]; 59; 65; 0.626110; GGGTTTA; 0.00409; 0.00190;
Sequence; YY1 [T00915]; 39; 42; 0.000000; ATGG; 0.26172; 0.20374;
Sequence; YY1 [T00915]; 56; 59; 0.000000; ATGG; 0.26172; 0.20374;
Sequence; ER-alpha [T00261]; 47; 51; 0.000000; GGTCA; 0.06543; 0.04373;
Sequence; RAR-alpha1 [T00719]; 45; 57; 12.025302; CGGGTCACGTGAT; 0.00008; 0.00003;
Sequence; E2F-1 [T01542]; 10; 17; 5.042045; GCGGGAGC; 0.00613; 0.04525;
Sequence; E2F-1 [T01542]; 27; 34; 9.028527; GCGGGGAC; 0.00920; 0.04410;
Sequence; E2F-1 [T01542]; 44; 51; 7.839654; GCGGGTCA; 0.01022; 0.01993;
Sequence; Pax-5 [T00070]; 25; 31; 1.537547; GGGCGGG; 0.02454; 0.30539;
Sequence; Pax-5 [T00070]; 42; 48; 1.537547; GGGCGGG; 0.02454; 0.30539;
Sequence; p53 [T00671]; 25; 31; 3.375208; GGGCGGG; 0.02454; 0.27033;
Sequence; p53 [T00671]; 42; 48; 3.375208; GGGCGGG; 0.02454; 0.27033;
Sequence; Sp1 [T00759]; 24; 33; 1.388285; AGGGCGGGGA; 0.00109; 0.05209;
Sequence; Sp1 [T00759]; 41; 50; 1.229271; GGGGCGGGTC; 0.00045; 0.02841;
Sequence; MAZ [T00490]; 38; 50; 14.398434; GATGGGGCGGGTC; 0.00034; 0.02641;
Sequence; TFII-I [T00824]; 13; 18; 11.337888; GGAGCG; 0.08179; 0.35791;
Sequence; TFII-I [T00824]; 31; 36; 9.512894; GGACGG; 0.24536; 0.31372;
Sequence; WT1 [T00899]; 10; 18; 11.111111; GCGGGAGCG; 0.00690; 0.25187;
Sequence; WT1 [T00899]; 27; 35; 11.111111; GCGGGGACG; 0.00690; 0.25187;
Sequence; GCF [T00320]; 16; 24; 4.846987; GCGCCGAGA; 0.00920; 0.03702;

**>LHFPL5 AA-sequence**
CGGACGGAGGAGGGGGCGGGGCTCTCGGGAGCCGTGAGCCGGGAAGAGGGAGACGGGCAGGGCGGCG

-- Factors predicted by PROMO in this sequence ----------------------
NAME; MATRIX_WIDTH;
GR-alpha [T00337]; 5
Pax-5 [T00070]; 7
p53 [T00671]; 7
TFII-I [T00824]; 6
STAT4 [T01577]; 6
c-Ets-1 [T00112]; 7
Elk-1 [T00250]; 9
RAR-beta:RXR-alpha [T05420]; 12
E2F-1 [T01542]; 8
Sp1 [T00759]; 10
ETF [T00270]; 11
WT1 [T00899]; 9

-- PROMO predictions detail ------------------------------------------

Sequence name; Factor name; Start position; End position; Dissimilarity; String; RE equally; RE query
Sequence; GR-alpha [T00337]; 5; 9; 8.281568; GGAGG; 0.26172; 1.11326;
Sequence; GR-alpha [T00337]; 8; 12; 8.281568; GGAGG; 0.26172; 1.11326;
Sequence; GR-alpha [T00337]; 44; 48; 0.207689; AGAGG; 0.26172; 0.30889;
Sequence; GR-alpha [T00337]; 56; 60; 8.073878; GCAGG; 0.26172; 0.41270;
Sequence; Pax-5 [T00070]; 13; 19; 1.537547; GGGCGGG; 0.02454; 0.45486;
Sequence; Pax-5 [T00070]; 18; 24; 4.007279; GGGCTCT; 0.03680; 0.04366;
Sequence; Pax-5 [T00070]; 54; 60; 0.000000; GGGCAGG; 0.03680; 0.30880;
Sequence; Pax-5 [T00070]; 59; 65; 9.552105; GGGCGGC; 0.04907; 0.31219;
Sequence; p53 [T00671]; 13; 19; 3.375208; GGGCGGG; 0.02454; 0.38186;
Sequence; p53 [T00671]; 18; 24; 8.537081; GGGCTCT; 0.00409; 0.00337;
Sequence; p53 [T00671]; 54; 60; 0.000000; GGGCAGG; 0.01227; 0.10507;
Sequence; p53 [T00671]; 59; 65; 6.188498; GGGCGGC; 0.02045; 0.15274;
Sequence; TFII-I [T00824]; 1; 6; 9.512894; GGACGG; 0.24536; 0.41929;
Sequence; TFII-I [T00824]; 8; 13; 11.337888; GGAGGG; 0.08179; 0.57113;
Sequence; TFII-I [T00824]; 48; 53; 11.337888; GGAGAC; 0.08179; 0.57113;
Sequence; STAT4 [T01577]; 41; 46; 4.411765; GGAAGA; 0.06543; 0.11248;
Sequence; c-Ets-1 [T00112]; 39; 45; 5.430224; CGGGAAG; 0.01227; 0.09913;
Sequence; Elk-1 [T00250]; 37; 45; 10.962309; GCCGGGAAG; 0.00895; 0.03318;
Sequence; RAR-beta:RXR-alpha [T05420]; 18; 29; 7.477995; GGGCTCTCGGGA; 0.00096; 0.00530;
Sequence; E2F-1 [T01542]; 15; 22; 10.022110; GCGGGGCT; 0.01125; 0.09694;
Sequence; Sp1 [T00759]; 12; 21; 0.000000; GGGGCGGGGC; 0.00006; 0.01932;
Sequence; ETF [T00270]; 6; 16; 8.876947; GAGGAGGGGGC; 0.00080; 0.02886;
Sequence; ETF [T00270]; 11; 21; 5.246906; GGGGGCGGGGC; 0.00096; 0.11002;
Sequence; WT1 [T00899]; 9; 17; 11.111111; GAGGGGGCG; 0.00690; 0.42272;

**>LHFPL5 VA-sequence**
CGGACGGGGGAGGGGGCGGGGCTCTCGGGAGCCGTGAGCCGGGAAGAGGGAGACGGGCAGGGCGGCG

-- Factors predicted by PROMO in this sequence ----------------------
NAME; MATRIX_WIDTH;
GR-alpha [T00337]; 5
Pax-5 [T00070]; 7
p53 [T00671]; 7
TFII-I [T00824]; 6
STAT4 [T01577]; 6
c-Ets-1 [T00112]; 7
Elk-1 [T00250]; 9
RAR-beta:RXR-alpha [T05420]; 12
E2F-1 [T01542]; 8
Sp1 [T00759]; 10
ETF [T00270]; 11
WT1 [T00899]; 9
MAZ [T00490]; 13

-- PROMO predictions detail ------------------------------------------

Sequence name; Factor name; Start position; End position; Dissimilarity; String; RE equally; RE query
Sequence; GR-alpha [T00337]; 8; 12; 8.281568; GGAGG; 0.26172; 1.08529;
Sequence; GR-alpha [T00337]; 44; 48; 0.207689; AGAGG; 0.26172; 0.26892;
Sequence; GR-alpha [T00337]; 56; 60; 8.073878; GCAGG; 0.26172; 0.40384;
Sequence; Pax-5 [T00070]; 13; 19; 1.537547; GGGCGGG; 0.02454; 0.51412;
Sequence; Pax-5 [T00070]; 18; 24; 4.007279; GGGCTCT; 0.03680; 0.04344;
Sequence; Pax-5 [T00070]; 54; 60; 0.000000; GGGCAGG; 0.03680; 0.33369;
Sequence; Pax-5 [T00070]; 59; 65; 9.552105; GGGCGGC; 0.04907; 0.33062;
Sequence; p53 [T00671]; 13; 19; 3.375208; GGGCGGG; 0.02454; 0.43155;
Sequence; p53 [T00671]; 18; 24; 8.537081; GGGCTCT; 0.00409; 0.00305;
Sequence; p53 [T00671]; 54; 60; 0.000000; GGGCAGG; 0.01227; 0.10568;
Sequence; p53 [T00671]; 59; 65; 6.188498; GGGCGGC; 0.02045; 0.16612;
Sequence; TFII-I [T00824]; 1; 6; 9.512894; GGACGG; 0.24536; 0.39677;
Sequence; TFII-I [T00824]; 8; 13; 11.337888; GGAGGG; 0.08179; 0.57397;
Sequence; TFII-I [T00824]; 48; 53; 11.337888; GGAGAC; 0.08179; 0.57397;
Sequence; STAT4 [T01577]; 41; 46; 4.411765; GGAAGA; 0.06543; 0.09803;
Sequence; c-Ets-1 [T00112]; 39; 45; 5.430224; CGGGAAG; 0.01227; 0.09067;
Sequence; Elk-1 [T00250]; 37; 45; 10.962309; GCCGGGAAG; 0.00895; 0.02963;
Sequence; RAR-beta:RXR-alpha [T05420]; 18; 29; 7.477995; GGGCTCTCGGGA; 0.00096; 0.00524;
Sequence; E2F-1 [T01542]; 15; 22; 10.022110; GCGGGGCT; 0.01125; 0.10980;
Sequence; Sp1 [T00759]; 12; 21; 0.000000; GGGGCGGGGC; 0.00006; 0.02365;
Sequence; ETF [T00270]; 6; 16; 7.870358; GGGGAGGGGGC; 0.00240; 0.17110;
Sequence; ETF [T00270]; 11; 21; 5.246906; GGGGGCGGGGC; 0.00096; 0.12929;
Sequence; WT1 [T00899]; 9; 17; 11.111111; GAGGGGGCG; 0.00690; 0.49200;
Sequence; MAZ [T00490]; 3; 15; 3.986869; ACGGGGGAGGGGG; 0.00020; 0.01584;

**References:**

#### 1) Xavier Messeguer, Ruth Escudero, Domènec Farré, Oscar Nuñez, Javier Martínez, M.Mar Albà. PROMO: detection of known transcription regulatory elements using species-tailored searches. Bioinformatics, 18, 2, 333-334, 2002.

#### 2) Domènec Farré, Romà Roset, Mario Huerta, José E. Adsuara, Llorenç Roselló, M.Mar Albà, Xavier Messeguer. Identification of patterns in biological sequences at the ALGGEN server: PROMO and MALGEN. Nucleic Acids Res, 31, 13, 3651-3653, 2003.
