## Supplementary material for "Genetic variations in G-Quadruplex forming sequences affect the transcription of human disease-related genes": Graphical Abstract

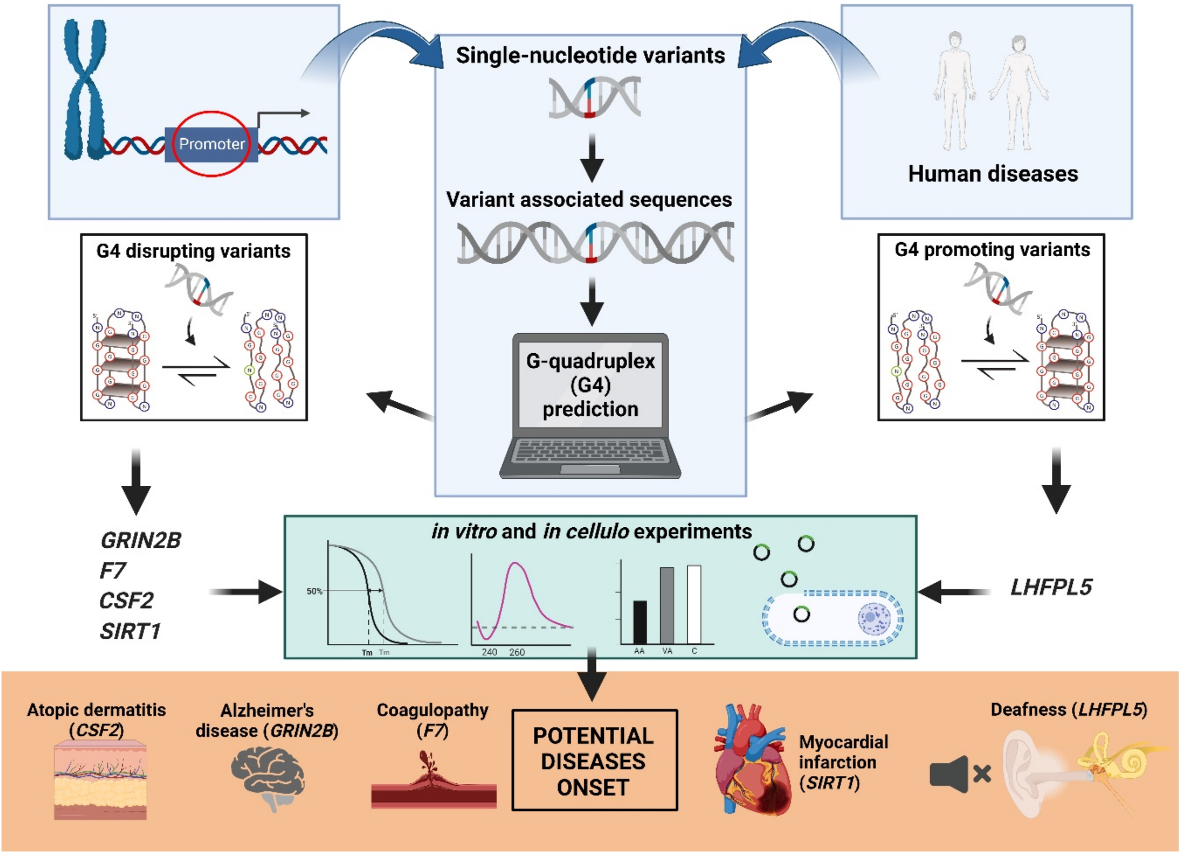


We have searched for human short genetic variants occurring in G-quadruplexes within promoters, demonstrating that they alter gene transcription and may be responsible for the onset of diseases (Created with BioRender.com).
